# DNA barcodes reveal inconsistent species boundaries in rose gall wasps *Diplolepis* and their inquilines *Periclistus* (Hymenoptera: Cynipidae)

**DOI:** 10.1101/530949

**Authors:** Y. Miles Zhang, Zoltán László, Chris Looney, Avar-Lehel Dénes, Robert H. Hanner, Joseph D. Shorthouse

## Abstract

Rose gall wasps *Diplolepis* induce structurally distinct galls on wild roses, which provide gallers with food and shelter. These galls are attacked by a wide variety of micro-hymenopterans including another cynipid *Periclistus* that act as inquilines. Both *Diplolepis* and *Periclistus* are difficult to distinguish based on adult morphology, instead the structural appearance of galls is often used to distinguish species. Using the mitochondrial gene *COI*, we test the species boundaries and built phylogenies of both *Diplolepis* and *Periclistus*. The molecular results have largely supported the validity of species described in the literature, with notable exceptions in four species groups. *Periclistus* exhibits a divide between the Palearctic and Nearctic clades, and ranges from specialists to generalists in terms of host specificity. While it is premature to enact any taxonomic changes without additional molecular markers, this incongruence between morphological and molecular data indicates these groups need taxonomic revision and gall morphology alone may be inadequate to delimit species.

## Introduction

Insect galls are one of the most spectacular products of evolution, as they represent atypical organ-like structures made by plants under the direction of stimuli provided by insects (Shorthouse et al. 2005). These novel plant structures provide food and protection from the elements for the galler (Ronquist et al. 2015; Stone et al. 2002). The ability to induce galls has evolved in seven orders of insects, but perhaps the most complex are those induced by cynipid wasps (Hymenoptera: Cynipidae). The majority of the approximately 1400 described species of gall wasps induce galls on leaves, stems, or roots of oaks (Fagaceae, *Quercus* L.) and roses (Rosaceae, *Rosa* L.) (Ronquist et al. 2015). Although seemingly well protected, cynipid galls attract many species of Hymenoptera with different feeding ecologies ranging from phytophagous inquilines that feed only on gall tissues, to parasitoids that feed on gall-inhabiting larvae (Hayward and Stone 2005). The assemblage of all inhabitants associated with a population of galls induced by the same gall wasp species is considered a component community, and each species of gall wasp is thought to support a unique gall community (Shorthouse 2010).

Interactions among and between cynipid species and their associated communities are complex, and construction of qualitative or quantitative food webs to understand these interactions is challenging (Stone et al. 2002). Many gall component communities are known to contain morphologically cryptic species (Abe et al. 2007; Nicholls et al. 2010; Nicholls et al. 2018; Zhang et al. 2014; Zhang et al. 2017), and the addition of molecular tools to aid in both species determination and the discovery of new species has helped resolve the complex relationships among cynipid gall component communities.

There have been approximately 50 species of gall wasps in the genus *Diplolepis* Geoffroy described worldwide, all of which inducing galls on roses. Nearly ⅔ of these were described from North America, suggesting the genus is poorly represented in the Palearctic. However, this could also reflect insufficient sampling, as evidenced by recent descriptions of new species from China (Wang et al. 2013). Species identification of *Diplolepis* based on adult morphology is challenging as few original descriptions have insufficient detail and identification keys are lacking (Shorthouse 1993; Shorthouse 2010). Phylogenetic relationships of *Diplolepis* have been investigated using two mitochondrial gene regions, cytochrome *b* and *12S* rRNA, but with conflicting results due to limited taxon sampling and poor sequence quality (Plantard et al. 1998).

Based on the results of their analyses, Plantard et al. (1998) divided the Nearctic *Diplolepis* species into four groups: “*nebulosa*”, “*polita*”, “*rosaefolii*”, and “flanged femur” clades. The Palearctic species were divided into two species groups: “*eglanteriae*” and “*rosae”*. However, the validity of *Diplolepis* species were not tested, despite some species differing by less than 10 base pairs out of 386 (0.026%) in *CytB* (Plantard et al. 1998). This brings further doubt into current identification of *Diplolepis*, which has traditionally separated species based on their distinctive galls.

Inquilines of the genus *Periclistus* Förster (Hymenoptera: Cynipidae) have lost the ability to induce their own galls (Ronquist et al. 2015), and are obligatorily dependent on completing their development within galls of *Diplolepis* (Brooks and Shorthouse 1998; Shorthouse and Brooks 1998). *Periclistus* induce gall tissues of their own from the tissues of the galls they inhabit. They do not feed on the bodies of the inducers, but the larval inducer is killed during oviposition by the female *Periclistus* (Shorthouse and Brooks 1998). Feeding by *Periclistus* causes each larva to be surrounded within its own chamber and as a result the inquiline-modified galls are structurally different from normal galls (Shorthouse 2010).

The phylogenetic position between inquilines and other gall-inducing cynipids has been controversial, ranging from a single origin of inquilism derived from gall-inducing cynipids (Liljeblad and Ronquist 1998) to multiple transitions between galler and inquilines (Ronquist et al. 2015). The genus *Periclistus* includes 18 described species worldwide, and all members of the genus are restricted to galls induced by *Diplolepis* to complete their larval development (Liljeblad and Ronquist 1998; Pujade-Villar et al. 2016; Ritchie 1984; Shorthouse and Brooks 1998). Ritchie (1984) revised the Nearctic *Periclistus* based on morphological characters in his PhD thesis, but the new species descriptions were not published and thus are not considered valid names.

DNA barcoding studies, which utilizes a 658 base pair region of the mitochondrial gene cytochrome *c* oxidase subunit I (*COI*) have demonstrated the ability of this marker to confidently link field collected organisms with a reference sequence of a previously identified species (Hebert et al. 2003). Species boundaries of *Diplolepis* and their associated inquiline *Periclistus* have been based exclusively on adult and gall morphology, but species identification is challenging in these genera (Ritchie 1984; Shorthouse 2010). Similar re-examination of species boundaries of cynipids and their associated parasitoids have demonstrated the utility of *COI* in integrative taxonomic revisions, which then leads to taxonomic revisions and description of new species (Ács et al. 2010; Zhang et al. 2014; Zhang et al. 2017). The major aim of this study is to: 1) test the species concepts of *Diplolepis*, and the inquiline *Periclistus* using *COI*; and 2) reconstruct the phylogeny of *Diplolepis* and *Periclistus*.

## Materials and Methods

### Specimen collection and deposition

The reference collection of coauthor JDS includes rose gall inhabitants collected over the past 50 years by himself and graduate students. Adults of *Diplolepis* and *Periclistus* were obtained by one of two ways. Mature galls initiated the previous year were collected in the spring after the inhabitants had been exposed to natural cold temperatures, storing them in either jars or whirl-pak bags at room temperature then removing the adults as they exited. Alternatively, mature galls were collected in the fall of the year they were induced, placed in whirl-pak bags and the galls subjected to temperatures of 0 to 3°C in incubators for 3 to 4 months to break diapause. The bags were then stored at room temperature and adults placed in alcohol as they exited the galls. In all cases, collections of the distinctive galls induced by each species were placed in separate bags or jars. This reference collection covers a wide geographical area across Canada, as well as representative collections from USA, Japan, and Turkey.

Representative specimens from all collection sites were pin-mounted. Reference collections of point-mounted specimens from many localities were deposited in the Canadian National Collection of Insects in Ottawa, Ontario; and the National Museum of Natural History in Washington D.C.. The remaining many hundreds of thousands of wet specimens were deposited at the University of Edinburgh in Edinburgh, Scotland under the care of Graham Stone. Additional voucher specimens from Northwestern USA were provided by coauthor CL, and are deposited at the Washington State Department of Agriculture Collection in Olympia, Washington. The Palearctic *Diplolepis* species used in this study were collected from Romania, Georgia, Russia, and Kazakhstan by coauthor ZL, and vouchers are stored in Babeş-Bolyai University, Cluj-Napoca, Romania.

All specimens used in this study were point mounted and identified to species whenever possible (see Table S1). Specimens of *Diplolepis* were identified by JDS (n=313), CL (n=14) or LZ (n=24). Specimens of *Periclistus* (n=260) were identified based on the unpublished key by Ritchie (1984). We opted to use numbers (e.g. *Periclistus* sp.1) to designate unnamed species as their species descriptions from Ritchie (1984)’s PhD dissertation are considered *nomina nuda* and invalid. The outgroups for the phylogenetic analyses of *Diplolepis* and *Periclistus* consisted of *Leibelia fukudae* (Shinji) and *Synophromorpha sylvestris* (Osten Sacken), respectively. The outgroups were chosen from published sequences of their closest relatives based on the phylogeny by Ronquist et al. (2015).

### DNA extraction and PCR amplification

The DNA extraction protocol was performed as part of a PhD thesis by Lima (2012). Genomic DNA was extracted from one or two legs removed from each voucher specimen using the methods outlined in Ivanova et al. (2006) at the Biodiversity Institute of Ontario, or at the Interdisciplinary Research Institute on Bio-Nano-Sciences of Babes-Bolyai University in Cluj-Napoca using the Qiagen Blood and Tissue Kit following standard protocol. The following primer sets were used to amplify the DNA barcode region of *COI*: LepF1 (5’-ATT CAA CCA ATC ATA AAG ATA TTG G-3’) and LepR1(5’-TAA ACT TCT GGA TGT CCA AAA AAT CA-3’) (Hebert et al. 2004); or MLepF1 (5’-GCT TTC CCA CGA ATA AAT A-3’) and MLepR1 (5’-CCT GTT CCA GCT CCA TTT TC-3’) (Hajibabaei et al. 2006); or LCO1490 (GGT CAA ATC ATA AAG ATA TTG G) and HCO2198 (TAA ACT TCA GGG TGA CCA AAA AAT CA) (Folmer et al. 1994).

PCR reactions were carried out in 96-well plates in 12.5 μL volumes containing: 2.5 mM MgCl2, 5 pmol of each primer, 20 mM dNTPs, 10 mM Tris-HCL (pH 8.3), 50 mM of KCl, 10-20 ng (1 to 2 μL) of genomic DNA and 1 unit *Taq* DNA polymerase (Platinum® Taq DNA polymerase, Invitrogen). PCR thermocycling profile was: 1 cycle of 60 seconds at 94°C, 5 cycles of 40 seconds at 94°C, 40 seconds at 45°C and 60 seconds at 72°C, followed by 35 cycles of 40 seconds at 94°C, at 51°C and 60 seconds at 72°C, with final extension of 5 minutes at 72°C. PCR products were visualized on a 2% agarose E-gel (Invitrogen), and positive single bands were selected for bi-directional sequencing with the BigDye Terminator Cycle Sequencing Kit on an ABI3730xl DNA Analyzer (Applied Biosystems) at the Biodiversity Institute of Ontario.

### Phylogenetic Analyses

Contigs of *COI* were assembled using Sequencher v4.5 (Gene Codes) and aligned using MUSCLE (Edgar 2004) implemented in MEGA X (Kumar et al. 2018) and manually inspected by eye for errors. Summary statistics were calculated using AMAS (Borowiec 2016). Bayesian inference analyses were conducted using MrBayes v3.2.6 (Ronquist et al. 2012). Each analysis had two independent searches with four chains and were run for 10,000,000 generations, sampling every 1000, with a 25% burnin discarded. The dataset was not partitioned based on nucleotide position and as it would limit the amount of data per partition (~219 bp) for accurate parameter estimation. The best fitting model of molecular evolution was tested using jModelTest2 (Darriba et al. 2012), and the general time-reversible model, with a parameter for invariant sites and rate heterogeneity modelled under a gamma distribution (GTR+I+G) was chosen based on the Bayesian Information Criterion (BIC) for both taxa. The phylogenetic trees were visualized in FigTree v1.4.2 (Rambaut 2012) and enhanced using Adobe Illustrator CS5. Intra- and interspecific divergence was calculated using the Kimura-2-Parameter (Kimura 1980) model in MEGA X. Automatic Barcode Gap Discovery (ABGD) was also performed using K2P model with default settings (Puillandre et al. 2012). Sequences are publicly available on GenBank (see Table S1), and the North American specimens can also be found on the Barcode of Life Data System (BOLD: http://barcodinglife.org/, Ratnasingham and Hebert 2007) in projects Rose gall wasps *Diplolepis* of North America (DIPNA) and Rose gall inquilines *Periclistus* of North America (PERNA).

## Results

### Gall Inducer *Diplolepis*

In total 313 *COI* sequences averaging 597bp (204 parsimony informative sites, 74% AT content) were generated from the *Diplolepis* specimens, including fifteen Nearctic and nine Palearctic species, as well as two undescribed species collected from Russia and Kazakhstan (Fig. 1, S1). The genus *Diplolepis* is recovered as monophyletic, and is further divided into the flanged femur and the leaf galler clades. Most the species were also recovered as monophyletic, with the exception of the following non-monophyletic sets of species among which genetic divergence was less than 3%: *D. polita* (Ashmead) and *D. bassetti* (Beutenmüller); *D. fusiformans* (Ashmead) *and D. rosaefolii* (Cockerell); *D. nebulosa* (Bassett), *D. variabilis* (Osten Sacken), and *D. ignota* (Bassett); *D. mayri* (Schlectendal), *D. rosae* (L.), *D. fructuum* (Rübsaamen) and *Diplolepis* sp.1. In contrast, round leaf galls resembling *D. japonica* (Walker) and *D. eglanteriae* (Hartig) collected in the Palearctic are genetically distinct from those collected in the Nearctic (3.43%), and were therefore split into two groups. The genetic distances are calculated by combining *Diplolepis* with very low divergences grouped into species groups, and intraspecific divergence ranged from 0 – 2.44% (Table S2A) while interspecific divergence ranged from 4.60 –17.55% (Table S3A).

**Figure 1.**
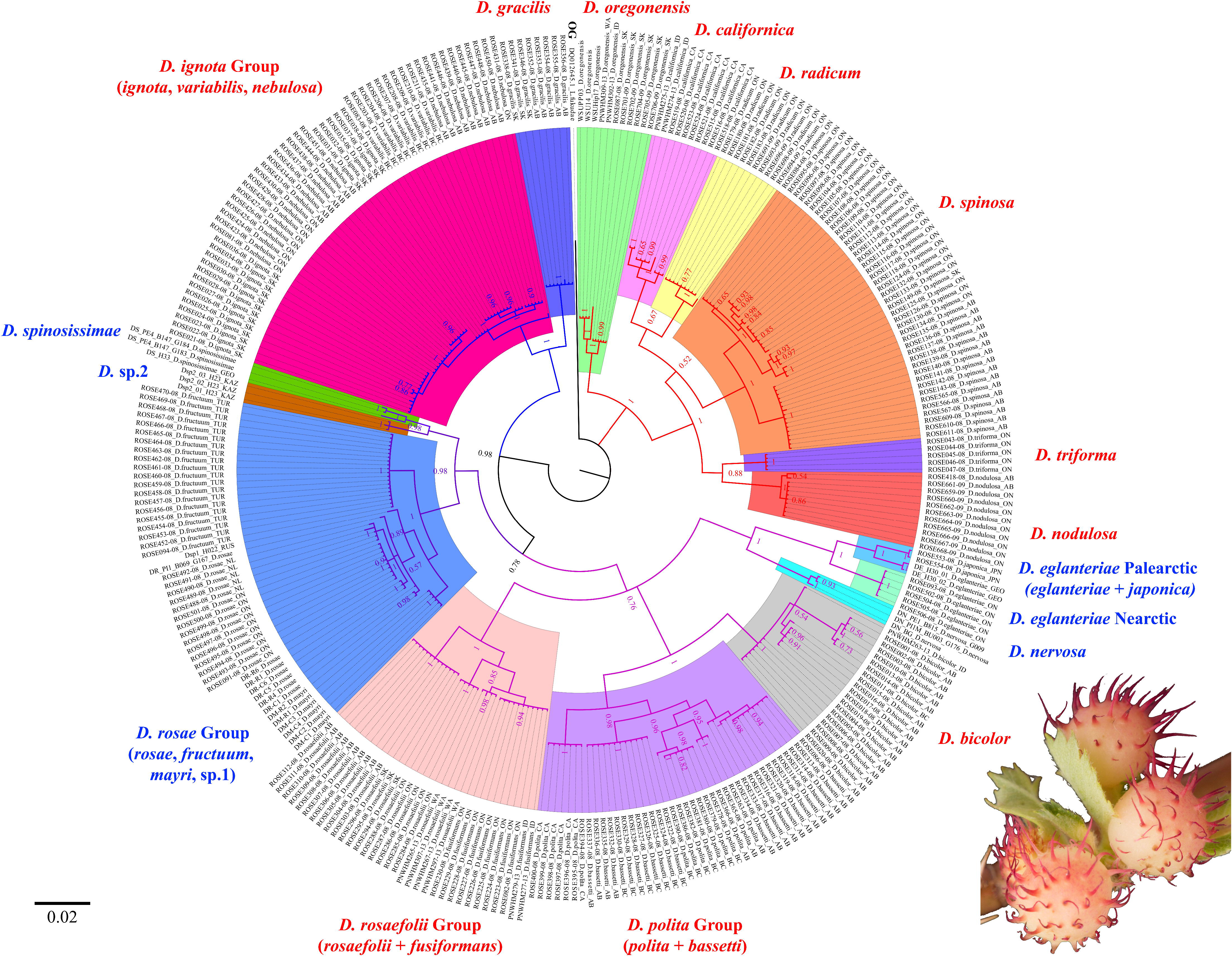
Bayesian inference tree of *Diplolepis* based on *COI*. Values indicate Bayesian posterior probability. The branches are colour coded by (sub)clades: red = Flanged femur, blue = Nearctic leaf galler, purple = Palearctic multi-chambered galler, and pink = Mixed leaf galler. Outgroup is labeled in black, while ingroup species are labelled as either Palearctic (Blue), Nearctic (Red), or OG (Outgroup). Photo of *D. polita* gall by YMZ.

### Inquiline *Periclistus*

A total of 260 *COI* sequences averaging 597bp (159 parsimony informative sites, 76% AT content) were used for the *Periclistus* analysis (Fig. 4, S2). The intraspecific divergence ranged from 0.13 – 2.16% (Table S2B) while interspecific divergence ranged from 3.55 – 9.30% (Table S3B). The genus *Periclistus* is recovered as monophyletic and is further divided into the Nearctic (*P. arefactus* McCracken and Egbert, *P. pirata* (Osten Sacken), *P. piceus* Fullaway, and two unidentified species labeled as *Periclistus* sp.1 and *Periclistus* sp.2) and Palearctic clades (*Periclistus brandtii* and *P. caninae*).

## Discussion

### Phylogeny of *Diplolepis* and Species Delimitation based on *COI*

This study is the first large molecular phylogenetic dataset to test the current species boundaries of the *Diplolepis* rose gall wasps. Most of the *Diplolepis* species were recovered as monophyletic groups, with the exception of *D. polita* + *D. bassetti*, *D. fusiformans* + *D. rosaefolii*, *D. ignota* + *D. nebulosa* + *D. variabilis*, and *D. mayri* + *D. rosae* + *D. fructuum* + *Diplolepis* sp.1 which were recovered with very little genetic distance between them (Fig. 1). In addition, the round leaf galls that resemble *D. eglanteriae*/*D. japonica* were split into Palearctic and Nearctic clades due to high genetic divergence. ABGD recovered a total of 17 species groups, which is consistent with our analysis (Fig. 1).

The *Diplolepis* tree divides into two major clades. The “flanged femur” clade was also recovered by Plantard et al. (1998), and includes exclusively Nearctic species that have the synapomorphic trait of flanged hind femora. Members of this clade oviposit on the stem tissue at the base of leaf buds, which develops as galls on stems [*D. triforma* Shorthouse & Ritchie, *D. californica* (Beutenmüller), *D. oregonensis* (Beutenmüller), *D. spinosa*, *D. nodulosa*] or adventitious roots [*D. radicum* (Osten Sacken)] of the plant.

The leaf galler clade includes all the species that do not have flanged hind femora, and includes species from both Palearctic and Nearctic regions that induce single or multi-chambered galls from either leaflets within buds or from tissues at the base of developing leaflets. Multiple species within the leaf galler clade have very low intraspecific genetic distance despite having distinct gall morphology (Fig. S1). This leaf galler clade was recovered as five separate lineages by Plantard et al. (1998), with the three Palearctic species grouping closer to the flanged femur clade. This polytomy observed by Plantard et al. (1998) is likely due to limited data, as they were only able to recover <400bp sequence fragments. This clade can be further split into three subclades. The Nearctic leaf galler subclade includes *D. gracilis* (Ashmead), *D. nebulosa*, *D. variabilis*, and *D. ignota*, and induces single or multi-chambered galls on leaves. The *ignota* group consists of *D. ignota*, *D. variabilis*, and *D. nebulosa*, all three of which induce spherical galls on the abaxial (lower) surface of leaves and have similar genetic sequences. Their galls range from single to multi-chambered, and are found on *R. arkansana* Porter (*D. ignota*) or *R. woodsii* (*D. variabilis* and *D. nebulosa*) from early spring to late summer (Shorthouse 2010). This result is congruent with Plantard et al. (1998), where only 1–3 base pair substitutions were observed in *CytB* between these three *Diplolepis* species.

The Palearctic multi-chamber subclade includes *D. fructuum*, *D. mayri*, *D. rosae*, *D. spinosissimae* (Giraud), and two undescribed species. *Diplolepis* sp.1 falls close to *D. rosae* with very little genetic divergence, but its gall may be single- or multi-chambered and appear on the leaf-vein or the stem. *Diplolepis* sp.2 is the sister group of *D. spinosissimae*, and induces single-chambered galls in the interior walls of hips. In our analysis the *rosae* group, which consists of *D. rosae*, *D. mayri*, *D. fructuum*, and *Diplolepis* sp.1 all have distinct gall morphology, but lack genetic variation based on *COI* data. In the past *D. fructuum* has been considered a geographic race of *D. mayri* (Güçlü et al. 2008), and our result once again casts doubt on the validity of these species. As *D. rosae* and *D. mayri* have been introduced to North America (Shorthouse 2001), we included samples of *D. rosae* from both its native and introduced range, which exhibited little genetic distance between populations.

Finally, there is a mixed leaf gall subclade including both Palearctic [*D. eglanteriae*, *D. japonica*, *D. nervosa* (Curtis)] and Nearctic species [*D. bicolor* (Harris), *D. polita*, *D. bassetti*, *D. rosaefolii*, *D. fusiformans*] (Güçlü et al. 2008; Shorthouse 2010). Almost all members of this group induce galls on leaf tissue, with the only exception being *D. fusiformans*, a species that forms small, fusiform galls on immature rose stems (Shorthouse 2010). The leaf galler *D. rosaefolii* was rendered paraphyletic by *D. fusiformans*. These two species are amongst the smallest Nearctic species, and are often found in the same habitat and on the same individual plant. It is possible that they are conspecific and capable of attacking both leaf and stem tissues. Similarly, the *polita* group consisting of *D. polita* and *D. bassetti* also have very little genetic distance, and both induce spiny, single-chambered galls on the adaxial (upper) surface of the leaf in the spring (Shorthouse 2010). The main differences between the two species are largely based on host plant and gall surface structures, as the galls of *D. polita* are generally weakly-spined and found on *R. acicularis* Lindl. and *R. nutkana* Presl. (Shorthouse 1973), whereas the galls induced by *D. bassetti* are mossy in appearance and mostly found on *R. woodsii* Lindl. (Shorthouse 2010). The Palearctic species *D. eglanteriae* was also thought to have been introduced to North America (Shorthouse 2001), however, specimens collected in Canada were genetically divergent from its conspecifics in the Palearctic. This is further confounded by the inclusion of *D. japonica* as the sister group to the Palearctic *D. eglanteriae* clade, which also induces round galls on rose leaves and is grouped together with the Palearctic *D. eglanteriae* clade. Therefore, we separated the round galls collected from the Palearctic and the Nearctic into two separate groups, but future studies with larger sample size from both Europe, Asia and North America is needed to fully delimit the boundaries of these species.

*Diplolepis* identification is primarily based on a combination of geography, host plant, and gall morphology rather than adult wasp morphology, which could have resulted in the over-splitting of species. Alternatively, mitochondrial genes such as *COI* and *CytB* may not delimit certain *Diplolepis* species complexes due to introgression or incomplete lineage sorting that leads to mitonuclear discordance, which has been observed in a variety of insects, including cynipids (Linnen and Farrell 2007; Nicholls et al. 2012; Rokas et al. 2003). Therefore, without the inclusion of additional nuclear genes and extensive morphological study of the type materials, we are hesitant to propose taxonomic changes based on *COI* data alone.

### Delimiting *Periclistus* using DNA barcodes

Similar to the gallers, the *COI* data were able to delimit the *Periclistus* species associated with *Diplolepis* galls into seven species (Fig. 2). *Periclistus caninae*, *P. brandtii*, *P. pirata*, *P. piceus*, and *Periclistus* sp.1 are inquilines of multiple species of galls: *P. caninae* and *P. pirata* attacks both single- and multi-chambered galls; *P. piceus* and *Periclistus* sp.1 reared exclusively from the single-chambered galls; while *P. brandtii* exclusively inhabits multi-chambered galls (Fig. S2). All five generalist *Periclistus* species are capable of modifying the small, single-chambered galls such as *D. nodulosa* and *D. polita* to larger, multi-chambered galls (Brooks and Shorthouse 1998; LeBlanc and Lacroix 2001; Shorthouse 1980). The presence of inquilines has been shown to change the community dynamics of the galls as the inducers are usually killed by *Periclistus* during oviposition (Shorthouse and Brooks 1998) and by altering the gall size and number of inhabitants in larger, multi-chambered galls where some inducers can survive the inquiline oviposition (László and Tóthmérész 2006). Additionally, this alteration in gall community also attracts additional specialist parasitoids that only feed on *Periclistus* (Zhang et al. 2014; Zhang et al. 2017). However, not all *Periclistus* attack multiple species of galls, as *P. arefactus* and *Periclistus* sp.2 are only associated with a single species of *Diplolepis*.

**Figure 2.**
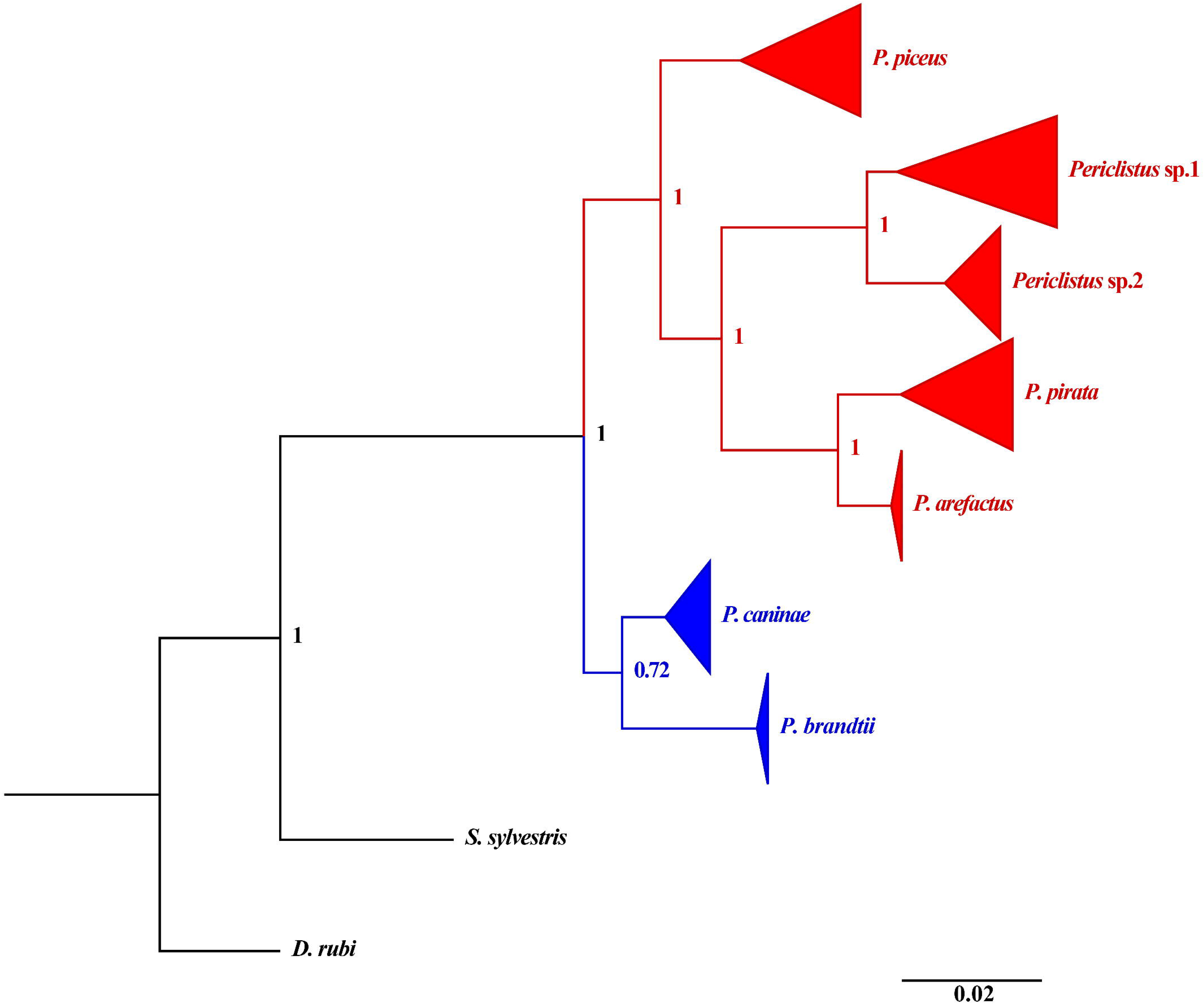
Bayesian inference tree of *Periclistus* based on *COI*. Values indicate Bayesian posterior probability. Outgroups are labeled in black, while ingroup species are labelled as either Palearctic (Blue) or Nearctic (Red).

With the addition of these two undescribed *Periclistus* species, the Holarctic diversity of *Periclistus* is increased to 14 species (Pujade-Villar et al. 2016). Our phylogeny includes less than half of the known species, so it is unclear whether this Palearctic/Nearctic divide will hold once more specimens are added. The description of these two new *Periclistus* species and the taxonomic revision of the genus are beyond the scope of this paper, however, we recommend revisions that uses molecular data as a guide for species descriptions as some of the morphological differences used by Ritchie (1984) differed from our *COI* results.

### Biogeography of rose, *Diplolepis*, and *Periclistus*

As is the case with most highly specialized phytophagous insects, *Diplolepis* gall-inducers are restricted to attacking closely related plants of the same genus. In the case of *Diplolepis* and *Periclistus*, all host plants are shrubs of the genus *Rosa* and the phylogeny of the insects cannot be understood without first discussing the host plants. Roses are notoriously difficult to identify, with some species are characterized by extensive continuous morphological variation that blurs their limits with each other and with their ancestors (Wissemann and Ritz 2007). Besides their intraspecific variability, hybridization and polyploidy resulting in species boundaries that are hard to define (Fougere-Danezan et al. 2015). However, the propensity to hybridize is likely a characteristic that provides new opportunities for *Diplolepis* to exploit and has contributed to speciation within the genus.

Based on recent biogeographic work on *Rosa*, the genus mostly likely evolved during Eocene in Asia and Western North America, and most extant American species are the results of re-colonization from Asia through the Bering Land Bridge (Fougere-Danezan et al. 2015). The genetic exchange between the two continents through the land bridge is also reflected in *Diplolepis* phylogeny, where multiple subclades within the leaf galler clade have mixed Palearctic and Nearctic species. The origin of *Diplolepis* is likely Palearctic, as the only fossil of the tribe Diplolepidini is found in Thorness Bay in United Kingdom which dates to Late Eocene (Antropov et al. 2014). This Palearctic origin is also strengthened by the fact that *Liebelia*, the sister group of *Diplolepis* that also attacks *Rosa*, is found exclusively in the Palearctic. Considering the high number of *Rosa* species in the Palearctic (>150), a larger number of undescribed *Diplolepis* species may be expected from Eastern Palearctic and Oriental region. Similar to the oak gall wasps Cynipini, which also have been historically under-studied in these two regions (Pénzes et al. 2018), sampling efforts are needed to fully understand the origin of Diplolepidini, and Cynipidae as a whole.

A similar evolutionary trend of increasing gall size is also observed in *Periclistus*, in which many species are able to modify single-chambered leaf galls into forming distinctly enlarged, multi-chambered galls (Brooks and Shorthouse 1998; LeBlanc and Lacroix 2001; Shorthouse 1980). All species of leaf gallers in North America are attacked by *Periclistus*; however, most of the stem galls are not (Shorthouse 2010), possibly suggesting that the ancestral *Periclistus* first attacked leaf galls of *Diplolepis*. Leaf galls are easily located and remain small and succulent for several weeks of their development, providing ample opportunity for ovipositing *Periclistus*. Once established in galls of one species, the resulting adults that exited galls late in the season could have oviposited in a different gall wasp species, setting the stage for sympatric speciation.

## Conclusion

The intimate relationships between gall wasps and their associated inquilines and parasitoids provides an ideal study system for evolutionary ecology and speciation. However, phylogenetic relationships in these groups remain unresolved. By using the *COI* marker in combination with wide sampling and detailed ecological data, we were able to build the largest phylogeny of the rose gall wasps *Diplolepis* to date. We also used the *COI* data to delimit species of *Diplolepis* and *Periclistus* and found disparity between gall morphology and molecular data. However, without additional genetic markers or morphological data of the wasps we chose not to propose taxonomic changes due to known biases of data interpretation based on a single mitochondrial gene. Regardless of the utilization of *COI* in cynipid phylogenetics, species identification based on gall or adult morphology should be viewed with caution, given the unresolved nature of these data. Future research should use an integrative taxonomic approach to resolving evolutionary relationships, and the incorporation of multi-locus or even genomic-level data should aid in the resolution of these cryptic but diverse groups of insects.

## Acknowledgements

We would like to acknowledge J. Lima for sequencing the Canadian *Diplolepis* and *Periclistus* specimens. Discovery Grants from the Natural Sciences and Engineering Research Council of Canada and the Laurentian University Research Fund supported the JDS field collections. The collecting trip of ZL to Russia, Kazakhstan and Georgia belonged to the RO-CRES expedition, funded by a project of the UBB Cluj, Romania with ID: CNFIS-FDI-2018-0104. Gall collections by CL in Washington State were partially supported by National Science Foundation-IGERT award no. 0114304.

---

**Supplementary Figure 1**. Full Bayesian inference tree of *Diplolepis* based on *COI*. Values indicate Bayesian posterior probability. The abbreviation after underscore in each specimen name indicates collecting location.

**Supplementary Figure 2.** Full Bayesian inference tree of *Periclistus* based on *COI*. Values indicate Bayesian posterior probability. The abbreviation after underscore in each specimen name indicates host *Diplolepis* gall.

**Supplementary Table 1.** Collection locality and GenBank accession numbers for all specimens used in this study.

**Supplementary Table 2.** Intraspecific genetic divergence of A) *Diplolepis* and B) *Periclistus*.

**Supplementary Table 3.** Interspecific genetic divergence of A) *Diplolepis* and B) *Periclistus*.

