## Supplementary figures and images for "DNA barcodes reveal inconsistent species boundaries in rose gall wasps *Diplolepis* and their inquilines *Periclistus* (Hymenoptera: Cynipidae)"

### Supplemental Figure 1

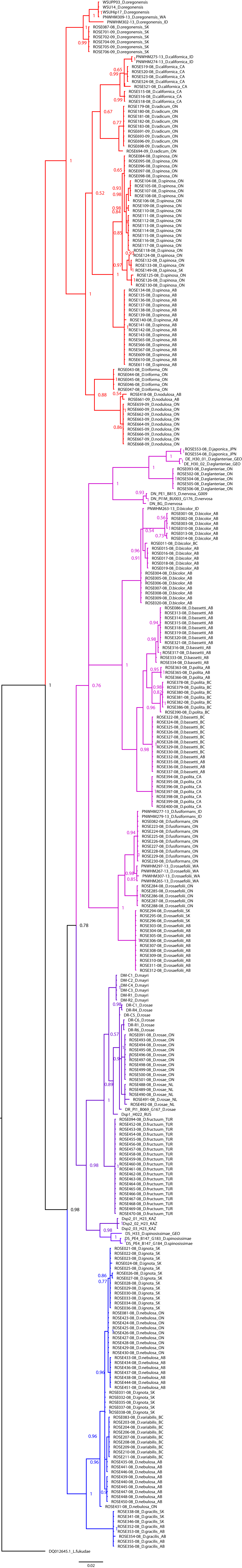

### Supplemental Figure 2

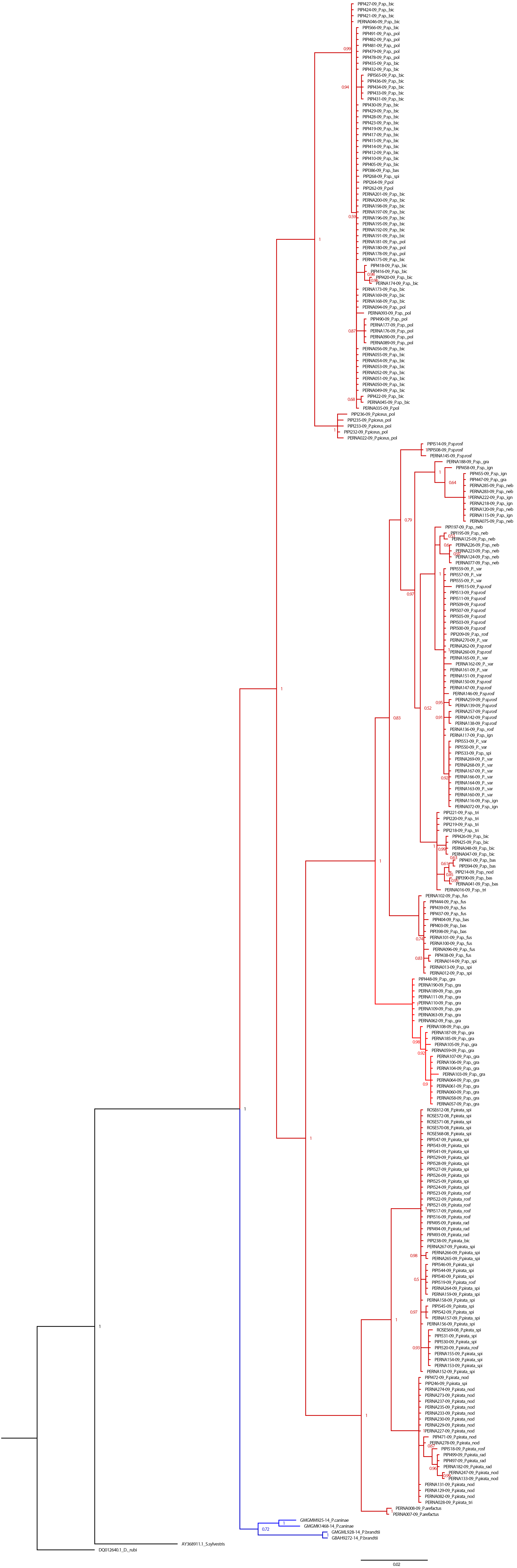
