## Supplemental Tables 1-3 for "DNA barcodes reveal inconsistent species boundaries in rose gall wasps *Diplolepis* and their inquilines *Periclistus* (Hymenoptera: Cynipidae)"

**Table S1.** Collection locality and GenBank accession numbers for all specimens used in this study.

| **Sample ID** | **Species** | **GenBank Accession** | **Country** | **Locality** | **Collection Date** | **GPS Coordinates** |
| --- | --- | --- | --- | --- | --- | --- |
| DE_H30_01 | D. eglanteriae | MK977672 | Georgia | Chartali | 13-Jul-2018 | 42.26, 44.68 |
| DE_H30_02 | D. eglanteriae | MK977673 | Georgia | Chartali | 13-Jul-2018 | 42.26, 44.68 |
| DM-C1 | D. mayri | MK977655 | Romania | Badesti | 12-Apr-2016 | 46.98, 23.63 |
| DM-C2 | D. mayri | MK977656 | Romania | Chidea | 07-Apr-2016 | 46.96, 23.59 |
| DM-C3 | D. mayri | MK977658 | Romania | Badesti | 13-Apr-2016 | 46.98, 23.63 |
| DM-C4 | D. mayri | MK977657 | Romania | Feiurdeni | 21-Mar-2016 | 46.9, 23.63 |
| DM-R1 | D. mayri | MK977659 | Romania | Luna de Jos | 01-Apr-2016 | 46.93, 23.72 |
| DM-R2 | D. mayri | MK977660 | Romania | Luna de Jos | 01-Apr-2016 | 46.92, 23.74 |
| DN_BG | D. nervosa | MK977675 | Romania | Cluj-Napoca | 31-Aug-2018 | 46.76, 23.59 |
| DN_PE1_B815 | D. nervosa | MK977674 | Romania | Chidea | 01-Aug-2018 | 46.96, 23.59 |
| DN_PI1M_BU003_G176 | D. nervosa | MK977676 | Romania | Cluj-Napoca | 01-Aug-2018 | 46.81, 23.65 |
| DR_PI1_B069_G167 | D. rosae | MK977677 | Romania | Cluj-Napoca | 01-Aug-2018 | 46.81, 23.65 |
| DR-C1 | D. rosae | MK977661 | Romania | Borsa-Catun | 07-Apr-2016 | 46.89, 23.71 |
| DR-C5 | D. rosae | MK977662 | Romania | Feiurdeni | 21-Mar-2016 | 46.9, 23.63 |
| DR-C6 | D. rosae | MK977663 | Romania | Borsa-Catun | 07-Apr-2016 | 46.89, 23.7 |
| DR-R1 | D. rosae | MK977664 | Romania | Feiurdeni | 21-Mar-2016 | 46.9, 23.63 |
| DR-R4 | D. rosae | MK977667 | Romania | Chidea | 07-Apr-2016 | 46.96, 23.59 |
| DR-R6 | D. rosae | MK977665 | Romania | Badesti | 13-Apr-2016 | 46.98, 23.63 |
| DS_H33 | D. spinosissimae | MK977671 | Georgia | Shorapani | 13-Jul-2018 | 42.09, 43.09 |
| DS_PE4_B147_G183 | D. spinosissimae | MK977678 | Romania | Cluj-Napoca | 01-Aug-2018 | 46.84, 23.62 |
| DS_PE4_B147_G184 | D. spinosissimae | MK977679 | Romania | Cluj-Napoca | 01-Aug-2018 | 46.84, 23.62 |
| Dsp1_H022 | D. sp.1 | MK977666 | Russia | Kozul'ka | 05-Jul-2018 | 56.18, 91.48 |
| Dsp2_01_H23 | D. sp.2 | MK977668 | Kazakhstan | Tavolzhan | 06-Jul-2018 | 52.71, 77.43 |
| Dsp2_02_H23 | D. sp.2 | MK977669 | Kazakhstan | Tavolzhan | 06-Jul-2018 | 52.71, 77.43 |
| Dsp2_03_H23 | D. sp.2 | MK977670 | Kazakhstan | Tavolzhan | 06-Jul-2018 | 52.71, 77.43 |
| PERNA007-09 | P. arefactus | HQ968038 | USA | Yolo Co. | 25-Apr-1967 | 38.49, -122.01 |
| PERNA008-09 | P. arefactus | HQ968039 | USA | Yolo Co. | 25-Apr-1967 | 38.49, -122.01 |
| PERNA012-09 | P. sp.1 | HQ968040 | Canada | Allumette Island | 04-May-1977 | 45.87, -77.17 |
| PERNA013-09 | P. sp.1 | HQ968041 | Canada | Arnprior | 02-May-1977 | 45.44, -76.36 |
| PERNA014-09 | P. sp.1 | HQ968042 | Canada | Arnprior | 02-May-1977 | 45.44, -76.36 |
| PERNA016-09 | P. sp.1 | HQ968043 | Canada | Manitoulin Island | 10-May-1979 | 45.95, -81.95 |
| PERNA022-09 | P. piceus | MK813814 | USA | Solano Co. | 01-Mar-1972 | 38.31, -121.9 |
| PERNA028-09 | P. pirata | MK813821 | Canada | Manitoulin Island | 07-May-1976 | 45.9, -82.23 |
| PERNA035-09 | P. piceus | MK813761 | Canada | Banff | 05-Oct-1981 | 51.18, -115.57 |
| PERNA041-09 | P. sp.1 | HQ968044 | Canada | Coaldale | 11-May-2002 | 49.87, -112.45 |
| PERNA045-09 | P. piceus | HQ968045 | Canada | Coaldale | 11-May-2003 | 49.87, -112.45 |
| PERNA046-09 | P. piceus | HQ968046 | Canada | Coaldale | 11-May-2003 | 49.87, -112.45 |
| PERNA047-09 | P. sp.1 | HQ968047 | Canada | Coaldale | 11-May-2003 | 49.87, -112.45 |
| PERNA048-09 | P. sp.1 | HQ968048 | Canada | Coaldale | 11-May-2003 | 49.87, -112.45 |
| PERNA049-09 | P. piceus | HQ968049 | Canada | Thunder Bay | 04-May-2005 | 48.38, -89.25 |
| PERNA050-09 | P. piceus | HQ968050 | Canada | Thunder Bay | 04-May-2005 | 48.38, -89.25 |
| PERNA051-09 | P. piceus | HQ968051 | Canada | Thunder Bay | 04-May-2005 | 48.38, -89.25 |
| PERNA052-09 | P. piceus | HQ968052 | Canada | Thunder Bay | 04-May-2005 | 48.38, -89.25 |
| PERNA053-09 | P. piceus | HQ968053 | Canada | Fort Albany | 27-May-2005 | 52.25, -81.5 |
| PERNA054-09 | P. piceus | HQ968054 | Canada | Fort Albany | 27-May-2005 | 52.25, -81.5 |
| PERNA055-09 | P. piceus | HQ968055 | Canada | Fort Albany | 27-May-2005 | 52.25, -81.5 |
| PERNA056-09 | P. piceus | HQ968056 | Canada | Fort Albany | 27-May-2005 | 52.25, -81.5 |
| PERNA057-09 | P. sp.2 | HQ968057 | Canada | Douglas P.P. | 26-Sep-1999 | 51.05, -106.46 |
| PERNA058-09 | P. sp.2 | HQ968058 | Canada | Douglas P.P. | 26-Sep-1999 | 51.05, -106.46 |
| PERNA059-09 | P. sp.2 | HQ968059 | Canada | Saskatoon | 29-Sep-1999 | 51.49, -106.41 |
| PERNA060-09 | P. sp.2 | HQ968060 | Canada | Saskatoon | 29-Sep-1999 | 51.49, -106.41 |
| PERNA061-09 | P. sp.2 | HQ968061 | Canada | Saskatoon | 29-Sep-1999 | 51.49, -106.41 |
| PERNA062-09 | P. sp.2 | HQ968062 | Canada | Douglas P.P. | 30-Sep-1999 | 51.05, -106.46 |
| PERNA063-09 | P. sp.2 | HQ968063 | Canada | Douglas P.P. | 30-Sep-1999 | 51.05, -106.46 |
| PERNA064-09 | P. sp.2 | HQ968064 | Canada | Douglas P.P. | 30-Sep-1999 | 51.05, -106.46 |
| PERNA072-09 | P. sp.1 | HQ968065 | Canada | Coaldale | 10-May-2007 | 49.87, -112.45 |
| PERNA075-09 | P. sp.1 | HQ968066 | Canada | Coaldale | 24-Oct-2002 | 49.87, -112.45 |
| PERNA077-09 | P. sp.1 | HQ968067 | Canada | Manitoulin Island | 28-Oct-2004 | 45.66, -82.26 |
| PERNA082-09 | P. sp.1 | HQ968068 | Canada | Timmins | 02-May-2002 | 48.48, -81.33 |
| PERNA089-09 | P. piceus | MK813762 | Canada | Winfield | 17-Oct-1999 | 50.04, -119.34 |
| PERNA090-09 | P. piceus | JN307250 | Canada | Winfield | 17-Oct-1999 | 50.04, -119.34 |
| PERNA093-09 | P. piceus | HQ968069 | Canada | Timmins | 11-Aug-2001 | 48.48, -81.33 |
| PERNA094-09 | P. piceus | HQ968070 | Canada | Timmins | 11-Aug-2001 | 48.48, -81.33 |
| PERNA096-09 | P. sp.1 | GU695012 | Canada | Renfrew | 28-Apr-2001 | 45.47, -76.69 |
| PERNA100-09 | P. sp.1 | GU695010 | Canada | Renfrew | 22-Apr-2003 | 45.47, -76.69 |
| PERNA101-09 | P. sp.1 | GU695011 | Canada | Renfrew | 22-Apr-2003 | 45.47, -76.69 |
| PERNA102-09 | P. sp.1 | GU695008 | Canada | Renfrew | 22-Apr-2003 | 45.47, -76.69 |
| PERNA103-09 | P. sp.2 | GU695009 | Canada | Renfrew | 22-Apr-2003 | 45.47, -76.69 |
| PERNA104-09 | P. sp.2 | GU695006 | Canada | Saskatoon | 29-Sep-1999 | 51.49, -106.41 |
| PERNA105-09 | P. sp.2 | GU695007 | Canada | Saskatoon | 29-Sep-1999 | 51.49, -106.41 |
| PERNA106-09 | P. sp.2 | GU695004 | Canada | Saskatoon | 29-Sep-1999 | 51.49, -106.41 |
| PERNA107-09 | P. sp.2 | GU695005 | Canada | Saskatoon | 29-Sep-1999 | 51.49, -106.41 |
| PERNA108-09 | P. sp.2 | GU695002 | Canada | Douglas P.P. | 30-Sep-1999 | 51.05, -106.46 |
| PERNA109-09 | P. sp.2 | GU695003 | Canada | Douglas P.P. | 30-Sep-1999 | 51.05, -106.46 |
| PERNA110-09 | P. sp.2 | GU695000 | Canada | Douglas P.P. | 30-Sep-1999 | 51.05, -106.46 |
| PERNA111-09 | P. sp.2 | GU695001 | Canada | Douglas P.P. | 30-Sep-1999 | 51.05, -106.46 |
| PERNA115-09 | P. sp.1 | GU694998 | Canada | Coaldale | 12-May-2003 | 49.87, -112.45 |
| PERNA116-09 | P. sp.1 | GU694995 | Canada | Coaldale | 10-May-2007 | 49.87, -112.45 |
| PERNA117-09 | P. sp.1 | GU694996 | Canada | Coaldale | 10-May-2007 | 49.87, -112.45 |
| PERNA120-09 | P. sp.1 | GU694992 | Canada | Coaldale | 24-Oct-2002 | 49.87, -112.45 |
| PERNA124-09 | P. sp.1 | GU694990 | Canada | Manitoulin Island | 28-Oct-2004 | 45.66, -82.26 |
| PERNA125-09 | P. sp.1 | GU694991 | Canada | Manitoulin Island | 28-Oct-2004 | 45.66, -82.26 |
| PERNA129-09 | P. pirata | GU694988 | Canada | Timmins | 02-May-2002 | 48.48, -81.33 |
| PERNA131-09 | P. pirata | GU694986 | Canada | Timmins | 02-May-2002 | 48.48, -81.33 |
| PERNA133-09 | P. pirata | GU694987 | Canada | Lethbridge | 09-May-2002 | 49.58, -112.83 |
| PERNA136-09 | P. sp.1 | GU694984 | Canada | Cochrane | 03-May-2002 | 49.06, -81.03 |
| PERNA138-09 | P. sp.1 | GU694985 | Canada | Cochrane | 03-May-2002 | 49.06, -81.03 |
| PERNA139-09 | P. sp.1 | GU694982 | Canada | Cochrane | 03-May-2002 | 49.06, -81.03 |
| PERNA142-09 | P. sp.1 | GU694983 | Canada | Cochrane | 03-May-2002 | 49.06, -81.03 |
| PERNA145-09 | P. sp.1 | GU694980 | Canada | Red Lake | 18-May-2002 | 51.02, -93.82 |
| PERNA146-09 | P. sp.1 | GU694981 | Canada | Red Lake | 18-May-2002 | 51.02, -93.82 |
| PERNA147-09 | P. sp.1 | GU694978 | Canada | Red Lake | 18-May-2002 | 51.02, -93.82 |
| PERNA150-09 | P. sp.1 | GU694979 | Canada | Dryden | 19-May-2002 | 49.78, -92.84 |
| PERNA151-09 | P. sp.1 | GU694976 | Canada | Dryden | 19-May-2002 | 49.78, -92.84 |
| PERNA152-09 | P. pirata | GU694977 | Canada | Waterton Lakes N.P. | 09-May-2007 | 49.08, -113.9 |
| PERNA153-09 | P. pirata | GU694974 | Canada | Waterton Lakes N.P. | 09-May-2007 | 49.08, -113.9 |
| PERNA154-09 | P. pirata | GU694975 | Canada | Waterton Lakes N.P. | 10-May-2007 | 49.08, -113.9 |
| PERNA155-09 | P. pirata | GU694972 | Canada | Waterton Lakes N.P. | 10-May-2007 | 49.08, -113.9 |
| PERNA156-09 | P. pirata | GU694973 | Canada | Waterton Lakes N.P. | 10-May-2007 | 49.08, -113.9 |
| PERNA157-09 | P. pirata | GU694970 | Canada | Waterton Lakes N.P. | 17-Sep-2007 | 49.08, -113.9 |
| PERNA158-09 | P. pirata | GU694971 | Canada | Waterton Lakes N.P. | 17-Sep-2007 | 49.08, -113.9 |
| PERNA159-09 | P. pirata | GU694968 | Canada | Waterton Lakes N.P. | 17-Sep-2007 | 49.08, -113.9 |
| PERNA160-09 | P. sp.1 | GU694969 | Canada | Peachland | 12-Feb-2008 | 49.79, -119.7 |
| PERNA161-09 | P. sp.1 | GU694966 | Canada | Peachland | 12-Feb-2008 | 49.79, -119.7 |
| PERNA162-09 | P. sp.1 | GU694967 | Canada | Peachland | 12-Feb-2008 | 49.79, -119.7 |
| PERNA163-09 | P. sp.1 | GU694964 | Canada | Peachland | 12-Feb-2008 | 49.79, -119.7 |
| PERNA164-09 | P. sp.1 | GU694965 | Canada | Peachland | 12-Feb-2008 | 49.79, -119.7 |
| PERNA165-09 | P. sp.1 | GU694962 | Canada | Peachland | 12-Feb-2008 | 49.79, -119.7 |
| PERNA166-09 | P. sp.1 | GU694963 | Canada | Peachland | 12-Feb-2008 | 49.79, -119.7 |
| PERNA167-09 | P. sp.1 | GU694960 | Canada | Peachland | 12-Feb-2008 | 49.79, -119.7 |
| PERNA168-09 | P. piceus | GU694961 | Canada | Coaldale | 11-May-2003 | 49.87, -112.45 |
| PERNA169-09 | P. piceus | GU694958 | Canada | Coaldale | 11-May-2003 | 49.87, -112.45 |
| PERNA173-09 | P. piceus | GU694956 | Canada | Fort Albany | 27-May-2005 | 52.25, -81.5 |
| PERNA174-09 | P. piceus | GU694957 | Canada | Fort Albany | 27-May-2005 | 52.25, -81.5 |
| PERNA175-09 | P. piceus | GU694954 | Canada | Fort Albany | 27-May-2005 | 52.25, -81.5 |
| PERNA176-09 | P. piceus | GU694955 | Canada | Winfield | 17-Oct-1999 | 50.04, -119.34 |
| PERNA177-09 | P. piceus | GU694952 | Canada | Winfield | 17-Oct-1999 | 50.04, -119.34 |
| PERNA178-09 | P. piceus | GU694953 | Canada | Timmins | 11-Aug-2001 | 48.48, -81.33 |
| PERNA180-09 | P. piceus | GU694950 | Canada | Timmins | 11-Aug-2001 | 48.48, -81.33 |
| PERNA181-09 | P. piceus | GU694951 | Canada | Timmins | 11-Aug-2001 | 48.48, -81.33 |
| PERNA182-09 | P. pirata | GU694948 | Canada | Liebenthal | 22-Sep-1999 | 50.66, -109.54 |
| PERNA185-09 | P. sp.2 | GU694949 | Canada | Saskatoon | 29-Sep-1999 | 51.49, -106.41 |
| PERNA187-09 | P. sp.2 | GU694947 | Canada | Saskatoon | 29-Sep-1999 | 51.49, -106.41 |
| PERNA188-09 | P. sp.1 | GU694945 | Canada | Douglas P.P. | 30-Sep-1999 | 51.05, -106.46 |
| PERNA189-09 | P. sp.2 | GU694946 | Canada | Douglas P.P. | 30-Sep-1999 | 51.05, -106.46 |
| PERNA190-09 | P. sp.2 | GU694944 | Canada | Douglas P.P. | 30-Sep-1999 | 51.05, -106.46 |
| PERNA191-09 | P. piceus | GU695151 | Canada | Coaldale | 11-May-2003 | 49.87, -112.45 |
| PERNA192-09 | P. piceus | GU695153 | Canada | Thunder Bay | 04-May-2005 | 48.38, -89.25 |
| PERNA195-09 | P. piceus | GU695155 | Canada | Thunder Bay | 04-May-2005 | 48.38, -89.25 |
| PERNA196-09 | P. piceus | GU695150 | Canada | Thunder Bay | 04-May-2005 | 48.38, -89.25 |
| PERNA197-09 | P. piceus | GU695148 | Canada | Fort Albany | 27-May-2005 | 52.25, -81.5 |
| PERNA198-09 | P. piceus | GU695149 | Canada | Fort Albany | 27-May-2005 | 52.25, -81.5 |
| PERNA200-09 | P. piceus | GU695154 | Canada | Fort Albany | 27-May-2005 | 52.25, -81.5 |
| PERNA201-09 | P. piceus | GU695152 | Canada | Fort Albany | 27-May-2005 | 52.25, -81.5 |
| PERNA218-09 | P. sp.1 | GU695156 | Canada | Coaldale | 12-May-2003 | 49.87, -112.45 |
| PERNA222-09 | P. sp.1 | GU695159 | Canada | Coaldale | 10-May-2007 | 49.87, -112.45 |
| PERNA223-09 | P. sp.1 | GU695160 | Canada | Manitoulin Island | 28-Oct-2004 | 45.66, -82.26 |
| PERNA226-09 | P. sp.1 | GU695161 | Canada | Manitoulin Island | 28-Oct-2004 | 45.66, -82.26 |
| PERNA227-09 | P. pirata | GU695141 | Canada | Timmins | 02-May-2002 | 48.48, -81.33 |
| PERNA229-09 | P. pirata | GU695137 | Canada | Timmins | 02-May-2002 | 48.48, -81.33 |
| PERNA230-09 | P. pirata | GU695140 | Canada | Timmins | 02-May-2002 | 48.48, -81.33 |
| PERNA233-09 | P. pirata | MK813833 | Canada | Timmins | 02-May-2002 | 48.48, -81.33 |
| PERNA235-09 | P. pirata | GU695139 | Canada | Timmins | 02-May-2002 | 48.48, -81.33 |
| PERNA237-09 | P. pirata | GU695134 | Canada | Timmins | 02-May-2002 | 48.48, -81.33 |
| PERNA247-09 | P. pirata | GU695143 | Canada | Lethbridge | 09-May-2002 | 49.58, -112.83 |
| PERNA257-09 | P. sp.1 | GU695168 | Canada | Cochrane | 03-May-2002 | 49.06, -81.03 |
| PERNA259-09 | P. sp.1 | GU695166 | Canada | Cochrane | 03-May-2002 | 49.06, -81.03 |
| PERNA260-09 | P. sp.1 | GU695164 | Canada | Cochrane | 03-May-2002 | 49.06, -81.03 |
| PERNA262-09 | P. sp.1 | GU695167 | Canada | Cochrane | 03-May-2002 | 49.06, -81.03 |
| PERNA264-09 | P. pirata | GU695145 | Canada | Waterton Lakes N.P. | 17-Sep-2007 | 49.08, -113.9 |
| PERNA265-09 | P. pirata | GU695147 | Canada | Waterton Lakes N.P. | 17-Sep-2007 | 49.08, -113.9 |
| PERNA266-09 | P. pirata | GU695144 | Canada | Waterton Lakes N.P. | 17-Sep-2007 | 49.08, -113.9 |
| PERNA267-09 | P. pirata | MK813835 | Canada | Waterton Lakes N.P. | 17-Sep-2007 | 49.08, -113.9 |
| PERNA268-09 | P. sp.1 | GU695162 | Canada | Peachland | 12-Feb-2008 | 49.79, -119.7 |
| PERNA269-09 | P. sp.1 | GU695163 | Canada | Peachland | 12-Feb-2008 | 49.79, -119.7 |
| PERNA270-09 | P. sp.1 | GU695165 | Canada | Peachland | 12-Feb-2008 | 49.79, -119.7 |
| PERNA273-09 | P. pirata | GU695138 | Canada | Timmins | 02-May-2002 | 48.48, -81.33 |
| PERNA274-09 | P. pirata | GU695135 | Canada | Timmins | 02-May-2002 | 48.48, -81.33 |
| PERNA278-09 | P. pirata | GU695142 | Canada | Renfrew | 23-Apr-2009 | 45.47, -76.69 |
| PERNA283-09 | P. sp.1 | GU695157 | Canada | Coaldale | 24-Oct-2002 | 49.87, -112.45 |
| PERNA285-09 | P. sp.1 | GU695158 | Canada | Coaldale | 24-Oct-2002 | 49.87, -112.45 |
| PIPI039-09 | P. sp.1 | MK813783 | Canada | Sudbury | 01-Sep-2008 | 46.47, -81 |
| PIPI160-09 | P. sp.1 | MK813784 | Canada | Sudbury | 01-Sep-2008 | 46.47, -81 |
| PIPI162-09 | P. sp.1 | MK813785 | Canada | Sudbury | 01-Sep-2008 | 46.47, -81 |
| PIPI163-09 | P. sp.1 | MK813786 | Canada | Sudbury | 01-Sep-2008 | 46.47, -81 |
| PIPI164-09 | P. sp.1 | MK813787 | Canada | Sudbury | 01-Sep-2008 | 46.47, -81 |
| PIPI165-09 | P. sp.1 | MK813788 | Canada | Sudbury | 01-Sep-2008 | 46.47, -81 |
| PIPI166-09 | P. sp.1 | MK813789 | Canada | Sudbury | 01-Sep-2008 | 46.47, -81 |
| PIPI168-09 | P. sp.1 | MK813790 | Canada | Sudbury | 01-Sep-2008 | 46.47, -81 |
| PIPI170-09 | P. sp.1 | MK813791 | Canada | Sudbury | 01-Sep-2008 | 46.47, -81 |
| PIPI171-09 | P. sp.1 | MK813792 | Canada | Sudbury | 01-Sep-2008 | 46.47, -81 |
| PIPI172-09 | P. sp.1 | MK813793 | Canada | Sudbury | 01-Sep-2008 | 46.47, -81 |
| PIPI173-09 | P. sp.1 | MK813794 | Canada | Sudbury | 01-Sep-2008 | 46.47, -81 |
| PIPI174-09 | P. sp.1 | MK813795 | Canada | Sudbury | 01-Sep-2008 | 46.47, -81 |
| PIPI175-09 | P. sp.1 | MK813796 | Canada | Sudbury | 01-Sep-2008 | 46.47, -81 |
| PIPI176-09 | P. sp.1 | MK813797 | Canada | Sudbury | 01-Sep-2008 | 46.47, -81 |
| PIPI177-09 | P. sp.1 | MK813798 | Canada | Sudbury | 01-Sep-2008 | 46.47, -81 |
| PIPI178-09 | P. sp.1 | MK813799 | Canada | Sudbury | 01-Sep-2008 | 46.47, -81 |
| PIPI195-09 | P. sp.1 | MK813801 | Canada | Manitoulin Island | 10-May-1979 | 45.66, -82.26 |
| PIPI197-09 | P. sp.1 | MK813800 | Canada | Manitoulin Island | 10-May-1979 | 45.66, -82.26 |
| PIPI209-09 | P. sp.1 | MK813802 | Canada | Sudbury | 25-May-1979 | 46.47, -81 |
| PIPI214-09 | P. sp.1 | MK813807 | Canada | Macklin | 15-Oct-1981 | 52.33, -109.94 |
| PIPI218-09 | P. sp.1 | MK813806 | Canada | Manitoulin Island | 10-May-1979 | 45.95, -81.95 |
| PIPI219-09 | P. sp.1 | MK813805 | Canada | Manitoulin Island | 10-May-1979 | 45.95, -81.95 |
| PIPI220-09 | P. sp.1 | MK813804 | Canada | Manitoulin Island | 10-May-1979 | 45.95, -81.95 |
| PIPI221-09 | P. sp.1 | MK813803 | Canada | Manitoulin Island | 10-May-1979 | 45.95, -81.95 |
| PIPI232-09 | P. piceus | MK813816 | USA | Solano Co. | 01-Mar-1972 | 38.31, -121.9 |
| PIPI233-09 | P. piceus | MK813813 | USA | Solano Co. | 01-Mar-1972 | 38.31, -121.9 |
| PIPI235-09 | P. piceus | MK813815 | USA | Solano Co. | 01-Mar-1972 | 38.31, -121.9 |
| PIPI236-09 | P. piceus | MK813812 | USA | Solano Co. | 01-Mar-1972 | 38.31, -121.9 |
| PIPI238-09 | P. pirata | MK813820 | Canada | Macklin | 15-Oct-1981 | 52.33, -109.94 |
| PIPI246-09 | P. pirata | MK813819 | Canada | Manitoulin Island | 02-May-1978 | 45.95, -81.95 |
| PIPI262-09 | P. piceus | MK813810 | Canada | Dawson city | 16-Aug-1971 | 64.04, -139.42 |
| PIPI264-09 | P. piceus | MK813809 | Canada | Banff | 05-Oct-1981 | 51.18, -115.57 |
| PIPI268-09 | P. piceus | MK813808 | Canada | Lethbridge | 12-Oct-1981 | 49.58, -112.83 |
| PIPI386-09 | P. piceus | GU695038 | Canada | Pincher Creek | 13-May-2003 | 49.49, -113.95 |
| PIPI390-09 | P. sp.1 | GU695039 | Canada | Coaldale | 11-May-2002 | 49.87, -112.45 |
| PIPI394-09 | P. sp.1 | GU695040 | Canada | Coaldale | 11-May-2002 | 49.87, -112.45 |
| PIPI398-09 | P. sp.1 | GU695042 | Canada | Coaldale | 24-Oct-2002 | 49.87, -112.45 |
| PIPI401-09 | P. sp.1 | GU695043 | Canada | Coaldale | 24-Oct-2002 | 49.87, -112.45 |
| PIPI403-09 | P. sp.1 | GU695044 | Canada | Coaldale | 24-Oct-2002 | 49.87, -112.45 |
| PIPI404-09 | P. sp.1 | GU695045 | Canada | Coaldale | 24-Oct-2002 | 49.87, -112.45 |
| PIPI405-09 | P. piceus | GU695046 | Canada | Thunder Bay | 04-May-2005 | 48.38, -89.25 |
| PIPI410-09 | P. piceus | GU695047 | Canada | Thunder Bay | 04-May-2005 | 48.38, -89.25 |
| PIPI412-09 | P. piceus | GU695048 | Canada | Thunder Bay | 04-May-2005 | 48.38, -89.25 |
| PIPI414-09 | P. piceus | GU695049 | Canada | Fort Albany | 27-May-2005 | 52.25, -81.5 |
| PIPI415-09 | P. piceus | GU695050 | Canada | Fort Albany | 27-May-2005 | 52.25, -81.5 |
| PIPI416-09 | P. piceus | GU695051 | Canada | Fort Albany | 27-May-2005 | 52.25, -81.5 |
| PIPI417-09 | P. piceus | GU695052 | Canada | Fort Albany | 27-May-2005 | 52.25, -81.5 |
| PIPI418-09 | P. piceus | GU695053 | Canada | Fort Albany | 27-May-2005 | 52.25, -81.5 |
| PIPI419-09 | P. piceus | GU695054 | Canada | Fort Albany | 27-May-2005 | 52.25, -81.5 |
| PIPI420-09 | P. piceus | GU695055 | Canada | Fort Albany | 27-May-2005 | 52.25, -81.5 |
| PIPI421-09 | P. piceus | GU695056 | Canada | Coaldale | 11-May-2003 | 49.87, -112.45 |
| PIPI422-09 | P. piceus | GU695057 | Canada | Coaldale | 11-May-2003 | 49.87, -112.45 |
| PIPI423-09 | P. piceus | GU695058 | Canada | Coaldale | 11-May-2003 | 49.87, -112.45 |
| PIPI424-09 | P. piceus | GU695059 | Canada | Coaldale | 11-May-2003 | 49.87, -112.45 |
| PIPI425-09 | P. sp.1 | GU695060 | Canada | Coaldale | 11-May-2003 | 49.87, -112.45 |
| PIPI426-09 | P. sp.1 | GU695061 | Canada | Coaldale | 11-May-2003 | 49.87, -112.45 |
| PIPI427-09 | P. piceus | GU695062 | Canada | Coaldale | 11-May-2003 | 49.87, -112.45 |
| PIPI428-09 | P. piceus | GU695063 | Canada | Coaldale | 11-May-2003 | 49.87, -112.45 |
| PIPI429-09 | P. piceus | GU695064 | Canada | Coaldale | 24-Oct-2002 | 49.87, -112.45 |
| PIPI430-09 | P. piceus | GU695065 | Canada | Coaldale | 24-Oct-2002 | 49.87, -112.45 |
| PIPI431-09 | P. piceus | GU695066 | Canada | Coaldale | 24-Oct-2002 | 49.87, -112.45 |
| PIPI432-09 | P. piceus | GU695067 | Canada | Coaldale | 24-Oct-2002 | 49.87, -112.45 |
| PIPI433-09 | P. piceus | GU695068 | Canada | Coaldale | 24-Oct-2002 | 49.87, -112.45 |
| PIPI434-09 | P. piceus | GU695069 | Canada | Coaldale | 24-Oct-2002 | 49.87, -112.45 |
| PIPI435-09 | P. piceus | GU695070 | Canada | Coaldale | 24-Oct-2002 | 49.87, -112.45 |
| PIPI436-09 | P. piceus | GU695071 | Canada | Coaldale | 24-Oct-2002 | 49.87, -112.45 |
| PIPI437-09 | P. sp.1 | GU695072 | Canada | Renfrew | 22-Apr-2003 | 45.47, -76.69 |
| PIPI438-09 | P. sp.1 | GU695073 | Canada | Renfrew | 22-Apr-2003 | 45.47, -76.69 |
| PIPI439-09 | P. sp.1 | GU695074 | Canada | Renfrew | 22-Apr-2003 | 45.47, -76.69 |
| PIPI444-09 | P. sp.1 | GU695075 | Canada | Renfrew | 28-Apr-2001 | 45.47, -76.69 |
| PIPI447-09 | P. sp.1 | GU695078 | Canada | Douglas P.P. | 26-Sep-1999 | 51.05, -106.46 |
| PIPI448-09 | P. sp.2 | GU695079 | Canada | Douglas P.P. | 26-Sep-1999 | 51.05, -106.46 |
| PIPI455-09 | P. sp.1 | GU695086 | Canada | Coaldale | 10-May-2007 | 49.87, -112.45 |
| PIPI458-09 | P. sp.1 | GU695087 | Canada | Coaldale | 10-May-2007 | 49.87, -112.45 |
| PIPI471-09 | P. pirata | GU695094 | Canada | Timmins | 02-May-2002 | 48.48, -81.33 |
| PIPI472-09 | P. pirata | GU695095 | Canada | Timmins | 02-May-2002 | 48.48, -81.33 |
| PIPI478-09 | P. piceus | GU695096 | Canada | Timmins | 11-Aug-2001 | 48.48, -81.33 |
| PIPI479-09 | P. piceus | GU695097 | Canada | Timmins | 11-Aug-2001 | 48.48, -81.33 |
| PIPI481-09 | P. piceus | MK813811 | Canada | Timmins | 11-Aug-2001 | 48.48, -81.33 |
| PIPI482-09 | P. piceus | GU695098 | Canada | Timmins | 11-Aug-2001 | 48.48, -81.33 |
| PIPI490-09 | P. piceus | GU695099 | Canada | Winfield | 17-Oct-1999 | 50.04, -119.34 |
| PIPI491-09 | P. piceus | GU695100 | Canada | Winfield | 17-Oct-1999 | 50.04, -119.34 |
| PIPI493-09 | P. pirata | GU695101 | Canada | Liebenthal | 22-Sep-1999 | 50.66, -109.54 |
| PIPI494-09 | P. pirata | GU695102 | Canada | Liebenthal | 22-Sep-1999 | 50.66, -109.54 |
| PIPI495-09 | P. pirata | GU695103 | Canada | Liebenthal | 22-Sep-1999 | 50.66, -109.54 |
| PIPI497-09 | P. pirata | GU695104 | Canada | Liebenthal | 22-Sep-1999 | 50.66, -109.54 |
| PIPI499-09 | P. pirata | GU695105 | Canada | Liebenthal | 22-Sep-1999 | 50.66, -109.54 |
| PIPI500-09 | P. sp.1 | GU695106 | Canada | Cochrane | 03-May-2002 | 49.06, -81.03 |
| PIPI503-09 | P. sp.1 | GU695107 | Canada | Cochrane | 03-May-2002 | 49.06, -81.03 |
| PIPI505-09 | P. sp.1 | GU695108 | Canada | Cochrane | 03-May-2002 | 49.06, -81.03 |
| PIPI507-09 | P. sp.1 | GU695109 | Canada | Cochrane | 03-May-2002 | 49.06, -81.03 |
| PIPI508-09 | P. sp.1 | GU695110 | Canada | Red Lake | 18-May-2002 | 51.02, -93.82 |
| PIPI509-09 | P. sp.1 | GU695111 | Canada | Red Lake | 18-May-2002 | 51.02, -93.82 |
| PIPI511-09 | P. sp.1 | GU695113 | Canada | Red Lake | 18-May-2002 | 51.02, -93.82 |
| PIPI513-09 | P. sp.1 | GU695115 | Canada | Red Lake | 18-May-2002 | 51.02, -93.82 |
| PIPI514-09 | P. sp.1 | GU695116 | Canada | Red Lake | 18-May-2002 | 51.02, -93.82 |
| PIPI515-09 | P. sp.1 | GU695117 | Canada | Red Lake | 18-May-2002 | 51.02, -93.82 |
| PIPI516-09 | P. pirata | GU695020 | Canada | Dryden | 19-May-2002 | 49.78, -92.84 |
| PIPI517-09 | P. pirata | GU695021 | Canada | Dryden | 19-May-2002 | 49.78, -92.84 |
| PIPI518-09 | P. pirata | GU695022 | Canada | Dryden | 19-May-2002 | 49.78, -92.84 |
| PIPI519-09 | P. pirata | GU695023 | Canada | Dryden | 19-May-2002 | 49.78, -92.84 |
| PIPI520-09 | P. pirata | GU695024 | Canada | Dryden | 19-May-2002 | 49.78, -92.84 |
| PIPI521-09 | P. pirata | GU695025 | Canada | Dryden | 19-May-2002 | 49.78, -92.84 |
| PIPI522-09 | P. pirata | GU695026 | Canada | Dryden | 19-May-2002 | 49.78, -92.84 |
| PIPI523-09 | P. pirata | GU695027 | Canada | Dryden | 19-May-2002 | 49.78, -92.84 |
| PIPI524-09 | P. pirata | GU695028 | Canada | Waterton Lakes N.P. | 09-May-2007 | 49.08, -113.9 |
| PIPI525-09 | P. pirata | GU695029 | Canada | Waterton Lakes N.P. | 09-May-2007 | 49.08, -113.9 |
| PIPI526-09 | P. pirata | GU695030 | Canada | Waterton Lakes N.P. | 09-May-2007 | 49.08, -113.9 |
| PIPI527-09 | P. pirata | GU695031 | Canada | Waterton Lakes N.P. | 09-May-2007 | 49.08, -113.9 |
| PIPI528-09 | P. pirata | GU695032 | Canada | Waterton Lakes N.P. | 09-May-2007 | 49.08, -113.9 |
| PIPI529-09 | P. pirata | GU695033 | Canada | Waterton Lakes N.P. | 09-May-2007 | 49.08, -113.9 |
| PIPI530-09 | P. pirata | GU695034 | Canada | Waterton Lakes N.P. | 09-May-2007 | 49.08, -113.9 |
| PIPI531-09 | P. pirata | GU695035 | Canada | Waterton Lakes N.P. | 09-May-2007 | 49.08, -113.9 |
| PIPI533-09 | P. sp.1 | GU695036 | Canada | Waterton Lakes N.P. | 10-May-2007 | 49.08, -113.9 |
| PIPI540-09 | P. pirata | GU695118 | Canada | Waterton Lakes N.P. | 17-Sep-2007 | 49.08, -113.9 |
| PIPI541-09 | P. pirata | GU695119 | Canada | Waterton Lakes N.P. | 17-Sep-2007 | 49.08, -113.9 |
| PIPI542-09 | P. pirata | GU695120 | Canada | Waterton Lakes N.P. | 17-Sep-2007 | 49.08, -113.9 |
| PIPI543-09 | P. pirata | GU695121 | Canada | Waterton Lakes N.P. | 17-Sep-2007 | 49.08, -113.9 |
| PIPI544-09 | P. pirata | GU695122 | Canada | Waterton Lakes N.P. | 17-Sep-2007 | 49.08, -113.9 |
| PIPI545-09 | P. pirata | GU695123 | Canada | Waterton Lakes N.P. | 17-Sep-2007 | 49.08, -113.9 |
| PIPI546-09 | P. pirata | GU695124 | Canada | Waterton Lakes N.P. | 17-Sep-2007 | 49.08, -113.9 |
| PIPI547-09 | P. pirata | GU695125 | Canada | Waterton Lakes N.P. | 17-Sep-2007 | 49.08, -113.9 |
| PIPI550-09 | P. sp.1 | GU695013 | Canada | Peachland | 16-May-2007 | 49.79, -119.7 |
| PIPI553-09 | P. sp.1 | GU695015 | Canada | Peachland | 16-May-2007 | 49.79, -119.7 |
| PIPI555-09 | P. sp.1 | GU695017 | Canada | Peachland | 16-May-2007 | 49.79, -119.7 |
| PIPI557-09 | P. sp.1 | GU695018 | Canada | Peachland | 12-Feb-2008 | 49.79, -119.7 |
| PIPI559-09 | P. sp.1 | GU695127 | Canada | Peachland | 12-Feb-2008 | 49.79, -119.7 |
| PIPI565-09 | P. piceus | GU695130 | Canada | Coaldale | 24-Oct-2002 | 49.87, -112.45 |
| PIPI566-09 | P. piceus | GU695131 | Canada | Coaldale | 24-Oct-2002 | 49.87, -112.45 |
| PNWHM263-13 | D. bicolor | MK977683 | USA | Latah | 01-May-2007 | 46.69, -116.95 |
| PNWHM265-13 | D. rosaefolii | MK977691 | USA | Whitman | 01-May-2007 | 46.7, -117.13 |
| PNWHM267-13 | D. rosaefolii | MK977689 | USA | Whitman | 01-May-2007 | 46.7, -117.13 |
| PNWHM274-13 | D. californica | MK977685 | USA | Latah | 01-May-2007 | 46.69, -116.95 |
| PNWHM275-13 | D. californica | MK977684 | USA | Latah | 01-May-2007 | 46.69, -116.95 |
| PNWHM277-13 | D. fusiformans | MK977686 | USA | Latah | 01-May-2007 | 46.66, -116.99 |
| PNWHM279-13 | D. fusiformans | MK977687 | USA | Latah | 01-May-2007 | 46.66, -116.99 |
| PNWHM297-13 | D. rosaefolii | MK977688 | USA | Whitman | 01-May-2007 | 46.7, -117.14 |
| PNWHM302-13 | D. oregonensis | MK977692 | USA | Latah | 01-May-2007 | 46.69, -116.95 |
| PNWHM307-13 | D. rosaefolii | MK977690 | USA | Whitman | 01-May-2007 | 46.7, -117.14 |
| PNWHM309-13 | D. oregonensis | MK977693 | USA | Whitman | 01-May-2007 | 46.7, -117.14 |
| ROSE001-08 | D. bicolor | MK813497 | Canada | Lethbridge | 18-Sep-1999 | 49.58, -112.83 |
| ROSE002-08 | D. bicolor | MK813496 | Canada | Lethbridge | 18-Sep-1999 | 49.58, -112.83 |
| ROSE003-08 | D. bicolor | MK813495 | Canada | Lethbridge | 18-Sep-1999 | 49.58, -112.83 |
| ROSE004-08 | D. bicolor | MK813494 | Canada | Waterton Lakes N.P. | 26-Sep-2007 | 49.08, -113.88 |
| ROSE005-08 | D. bicolor | MK813493 | Canada | Waterton Lakes N.P. | 26-Sep-2007 | 49.08, -113.88 |
| ROSE006-08 | D. bicolor | MK813492 | Canada | Waterton Lakes N.P. | 26-Sep-2007 | 49.08, -113.88 |
| ROSE007-08 | D. bicolor | MK813491 | Canada | Waterton Lakes N.P. | 26-Sep-2007 | 49.08, -113.88 |
| ROSE008-08 | D. bicolor | MK813490 | Canada | Waterton Lakes N.P. | 26-Sep-2007 | 49.08, -113.88 |
| ROSE009-08 | D. bicolor | MK813489 | Canada | Waterton Lakes N.P. | 26-Sep-2007 | 49.08, -113.88 |
| ROSE010-08 | D. bicolor | MK813488 | Canada | Waterton Lakes N.P. | 26-Sep-2007 | 49.08, -113.88 |
| ROSE011-08 | D. bicolor | MK813487 | Canada | Kelowna | 15-Oct-1999 | 49.89, -119.5 |
| ROSE013-08 | D. bicolor | MK813486 | Canada | Coaldale | 24-Oct-2002 | 49.87, -112.45 |
| ROSE014-08 | D. bicolor | MK813485 | Canada | Coaldale | 24-Oct-2002 | 49.87, -112.45 |
| ROSE015-08 | D. bicolor | MK813484 | Canada | Coaldale | 11-May-2003 | 49.87, -112.45 |
| ROSE016-08 | D. bicolor | MK813483 | Canada | Coaldale | 11-May-2003 | 49.87, -112.45 |
| ROSE017-08 | D. bicolor | MK813482 | Canada | Pincher Creek | 13-May-2003 | 49.49, -113.95 |
| ROSE018-08 | D. bicolor | MK813481 | Canada | Pincher Creek | 13-May-2003 | 49.49, -113.95 |
| ROSE019-08 | D. bicolor | MK813480 | Canada | Pincher Creek | 13-May-2003 | 49.49, -113.95 |
| ROSE020-08 | D. bicolor | MK813498 | Canada | Waterton Lakes N.P. | 26-Sep-2007 | 49.08, -113.88 |
| ROSE021-08 | D. ignota | MK813562 | Canada | Near Saskatoon | 12-May-2007 | 51.89, -106.81 |
| ROSE022-08 | D. ignota | MK813561 | Canada | Near Saskatoon | 12-May-2007 | 51.89, -106.81 |
| ROSE023-08 | D. ignota | MK813563 | Canada | Near Saskatoon | 12-May-2007 | 51.89, -106.81 |
| ROSE024-08 | D. ignota | MK813564 | Canada | Near Saskatoon | 12-May-2007 | 51.89, -106.81 |
| ROSE025-08 | D. ignota | MK813565 | Canada | Near Saskatoon | 12-May-2007 | 51.89, -106.81 |
| ROSE026-08 | D. ignota | MK813566 | Canada | Near Saskatoon | 12-May-2007 | 51.89, -106.81 |
| ROSE027-08 | D. ignota | MK813567 | Canada | Near Saskatoon | 12-May-2007 | 51.89, -106.81 |
| ROSE028-08 | D. ignota | MK813568 | Canada | Near Saskatoon | 12-May-2007 | 51.89, -106.81 |
| ROSE029-08 | D. ignota | MK813569 | Canada | Near Saskatoon | 12-May-2007 | 51.89, -106.81 |
| ROSE030-08 | D. ignota | MK813570 | Canada | Near Saskatoon | 12-May-2007 | 51.89, -106.81 |
| ROSE031-08 | D. ignota | MK813571 | Canada | Near Saskatoon | 12-May-2007 | 51.89, -106.81 |
| ROSE032-08 | D. ignota | MK813572 | Canada | Near Saskatoon | 12-May-2007 | 51.89, -106.81 |
| ROSE033-08 | D. ignota | MK813573 | Canada | Near Saskatoon | 12-May-2007 | 51.89, -106.81 |
| ROSE034-08 | D. ignota | MK813574 | Canada | Near Saskatoon | 12-May-2007 | 51.89, -106.81 |
| ROSE035-08 | D. ignota | MK813575 | Canada | Near Saskatoon | 12-May-2007 | 51.89, -106.81 |
| ROSE036-08 | D. ignota | MK813576 | Canada | Near Saskatoon | 12-May-2007 | 51.89, -106.81 |
| ROSE037-08 | D. ignota | MK813559 | Canada | Near Saskatoon | 12-May-2007 | 51.89, -106.81 |
| ROSE038-08 | D. ignota | MK813560 | Canada | Near Saskatoon | 12-May-2007 | 51.89, -106.81 |
| ROSE043-08 | D. triforma | MK813737 | Canada | Sudbury | 12-Jan-2008 | 46.47, -81 |
| ROSE044-08 | D. triforma | MK813738 | Canada | Sudbury | 12-Jan-2008 | 46.47, -81 |
| ROSE045-08 | D. triforma | MK813729 | Canada | Sudbury | 12-Jan-2008 | 46.47, -81 |
| ROSE046-08 | D. triforma | MK813739 | Canada | Sudbury | 12-Jan-2008 | 46.47, -81 |
| ROSE047-08 | D. triforma | MK813730 | Canada | Sudbury | 12-Jan-2008 | 46.47, -81 |
| ROSE081-08 | D. nebulosa | MK813586 | Canada | Manitoulin Island | 28-Oct-2004 | 45.66, -82.26 |
| ROSE082-08 | D. fusiformans | MK813549 | Canada | Renfrew | 19-Apr-2006 | 45.5, -76.58 |
| ROSE083-08 | D. variabilis | MK813752 | Canada | Peachland | 16-May-2007 | 49.79, -119.7 |
| ROSE084-08 | D. spinosa | MK813725 | Canada | Renfrew | 19-Apr-2006 | 45.47, -76.58 |
| ROSE086-08 | D. bassetti | MK813475 | Canada | Coaldale | 24-Oct-2002 | 49.87, -112.43 |
| ROSE087-08 | D. oregonensis | MK813623 | Canada | Great Sand Hills | 09-May-2003 | 50.69, -109.27 |
| ROSE091-08 | D. rosae | MK813632 | Canada | Picton | 28-Apr-2003 | 43.95, -76.85 |
| ROSE093-08 | D. eglanteriae | MK813515 | Canada | Niagara Falls | 06-May-1996 | 43.09, -79.09 |
| ROSE094-08 | D. fructuum | MK813533 | Turkey | Kelkit | 28-Jan-2008 | 40.09, 39.56 |
| ROSE095-08 | D. spinosa | MK813724 | Canada | Renfrew | 19-Apr-2006 | 45.47, -76.58 |
| ROSE096-08 | D. spinosa | MK813723 | Canada | Renfrew | 19-Apr-2006 | 45.47, -76.58 |
| ROSE097-08 | D. spinosa | MK813722 | Canada | Renfrew | 19-Apr-2006 | 45.47, -76.58 |
| ROSE098-08 | D. spinosa | MK813721 | Canada | Renfrew | 19-Apr-2006 | 45.47, -76.58 |
| ROSE104-08 | D. spinosa | MK813720 | Canada | Thunder Bay | 04-May-2005 | 48.25, -89.7 |
| ROSE105-08 | D. spinosa | MK813719 | Canada | Thunder Bay | 04-May-2005 | 48.25, -89.7 |
| ROSE106-08 | D. spinosa | MK813718 | Canada | Thunder Bay | 04-May-2005 | 48.25, -89.7 |
| ROSE107-08 | D. spinosa | MK813717 | Canada | Thunder Bay | 04-May-2005 | 48.25, -89.7 |
| ROSE108-08 | D. spinosa | MK813716 | Canada | Thunder Bay | 04-May-2005 | 48.25, -89.7 |
| ROSE109-08 | D. spinosa | MK813715 | Canada | Attawapiskat | 25-May-2005 | 52.92, -82.44 |
| ROSE110-08 | D. spinosa | MK813714 | Canada | Attawapiskat | 25-May-2005 | 52.92, -82.44 |
| ROSE111-08 | D. spinosa | MK813713 | Canada | Attawapiskat | 25-May-2005 | 52.92, -82.44 |
| ROSE112-08 | D. spinosa | MK813712 | Canada | Attawapiskat | 25-May-2005 | 52.92, -82.44 |
| ROSE113-08 | D. spinosa | MK813711 | Canada | Attawapiskat | 25-May-2005 | 52.92, -82.44 |
| ROSE114-08 | D. spinosa | MK813710 | Canada | Fort Albany | 27-May-2005 | 52.21, -81.69 |
| ROSE115-08 | D. spinosa | MK813709 | Canada | Fort Albany | 27-May-2005 | 52.21, -81.69 |
| ROSE116-08 | D. spinosa | MK813708 | Canada | Fort Albany | 27-May-2005 | 52.21, -81.69 |
| ROSE117-08 | D. spinosa | MK813707 | Canada | Fort Albany | 27-May-2005 | 52.21, -81.69 |
| ROSE118-08 | D. spinosa | MK813706 | Canada | Fort Albany | 27-May-2005 | 52.21, -81.69 |
| ROSE124-08 | D. spinosa | MK813705 | Canada | Renfrew | 19-Apr-2006 | 45.31, -75.92 |
| ROSE125-08 | D. spinosa | MK813704 | Canada | Renfrew | 19-Apr-2006 | 45.31, -75.92 |
| ROSE126-08 | D. spinosa | MK813703 | Canada | Renfrew | 19-Apr-2006 | 45.31, -75.92 |
| ROSE130-08 | D. spinosa | MK813702 | Canada | Renfrew | 19-Apr-2006 | 45.31, -75.92 |
| ROSE132-08 | D. spinosa | MK813701 | Canada | Kanata | 19-Apr-2006 | 45.35, -75.79 |
| ROSE133-08 | D. spinosa | MK813700 | Canada | Kanata | 19-Apr-2006 | 45.35, -75.79 |
| ROSE134-08 | D. spinosa | MK813699 | Canada | Waterton Lakes N.P. | 10-May-2007 | 49.08, -113.88 |
| ROSE135-08 | D. spinosa | MK813698 | Canada | Waterton Lakes N.P. | 10-May-2007 | 49.08, -113.88 |
| ROSE136-08 | D. spinosa | MK813697 | Canada | Waterton Lakes N.P. | 10-May-2007 | 49.08, -113.88 |
| ROSE137-08 | D. spinosa | MK813696 | Canada | Waterton Lakes N.P. | 10-May-2007 | 49.08, -113.88 |
| ROSE138-08 | D. spinosa | MK813695 | Canada | Waterton Lakes N.P. | 10-May-2007 | 49.08, -113.88 |
| ROSE139-08 | D. spinosa | MK813694 | Canada | Waterton Lakes N.P. | 10-May-2007 | 49.08, -113.88 |
| ROSE140-08 | D. spinosa | MK813693 | Canada | Waterton Lakes N.P. | 10-May-2007 | 49.08, -113.88 |
| ROSE141-08 | D. spinosa | MK813692 | Canada | Waterton Lakes N.P. | 10-May-2007 | 49.08, -113.88 |
| ROSE142-08 | D. spinosa | MK813691 | Canada | Waterton Lakes N.P. | 10-May-2007 | 49.08, -113.88 |
| ROSE143-08 | D. spinosa | MK813690 | Canada | Waterton Lakes N.P. | 10-May-2007 | 49.08, -113.88 |
| ROSE149-08 | D. spinosa | MK813689 | Canada | Cypress Hills | 21-Sep-1999 | 49.46, -109.36 |
| ROSE179-08 | D. radicum | MK813624 | Canada | Fort Albany | 28-May-2005 | 52.11, -81.92 |
| ROSE180-08 | D. radicum | MK813625 | Canada | Fort Albany | 28-May-2005 | 52.11, -81.92 |
| ROSE181-08 | D. radicum | MK813626 | Canada | Fort Albany | 28-May-2005 | 52.11, -81.92 |
| ROSE182-08 | D. radicum | MK813627 | Canada | Fort Albany | 28-May-2005 | 52.11, -81.92 |
| ROSE183-08 | D. radicum | MK813628 | Canada | Fort Albany | 28-May-2005 | 52.11, -81.92 |
| ROSE203-08 | D. variabilis | MK813753 | Canada | Peachland | 16-May-2007 | 49.79, -119.7 |
| ROSE204-08 | D. variabilis | MK813754 | Canada | Peachland | 16-May-2007 | 49.79, -119.7 |
| ROSE206-08 | D. variabilis | MK813755 | Canada | Peachland | 16-May-2007 | 49.79, -119.7 |
| ROSE207-08 | D. variabilis | MK813756 | Canada | Peachland | 16-May-2007 | 49.79, -119.7 |
| ROSE208-08 | D. variabilis | MK813757 | Canada | Peachland | 16-May-2007 | 49.79, -119.7 |
| ROSE209-08 | D. variabilis | MK813758 | Canada | Peachland | 16-May-2007 | 49.79, -119.7 |
| ROSE210-08 | D. variabilis | MK813760 | Canada | Peachland | 16-May-2007 | 49.79, -119.7 |
| ROSE211-08 | D. variabilis | MK813759 | Canada | Peachland | 16-May-2007 | 49.79, -119.7 |
| ROSE223-08 | D. fusiformans | MK813536 | Canada | Renfrew | 19-Apr-2006 | 45.5, -76.58 |
| ROSE224-08 | D. fusiformans | MK813537 | Canada | Renfrew | 19-Apr-2006 | 45.5, -76.58 |
| ROSE225-08 | D. fusiformans | MK813540 | Canada | Renfrew | 19-Apr-2006 | 45.5, -76.58 |
| ROSE226-08 | D. fusiformans | MK813541 | Canada | Renfrew | 19-Apr-2006 | 45.5, -76.58 |
| ROSE227-08 | D. fusiformans | MK813542 | Canada | Renfrew | 19-Apr-2006 | 45.5, -76.58 |
| ROSE228-08 | D. fusiformans | MK813545 | Canada | Renfrew | 19-Apr-2006 | 45.5, -76.58 |
| ROSE229-08 | D. fusiformans | MK813544 | Canada | Renfrew | 19-Apr-2006 | 45.5, -76.58 |
| ROSE230-08 | D. fusiformans | MK813543 | Canada | Renfrew | 19-Apr-2006 | 45.5, -76.58 |
| ROSE284-08 | D. rosaefolii | MK813660 | Canada | Timmins | 04-Sep-1991 | 48.47, -81.23 |
| ROSE285-08 | D. rosaefolii | MK813661 | Canada | Timmins | 04-Sep-1991 | 48.47, -81.23 |
| ROSE286-08 | D. rosaefolii | MK813662 | Canada | Timmins | 04-Sep-1991 | 48.47, -81.23 |
| ROSE287-08 | D. rosaefolii | MK813663 | Canada | Timmins | 04-Sep-1991 | 48.47, -81.23 |
| ROSE288-08 | D. rosaefolii | MK813664 | Canada | Timmins | 04-Sep-1991 | 48.47, -81.23 |
| ROSE294-08 | D. rosaefolii | MK813665 | Canada | Douglas P.P. | 26-Sep-1999 | 51.04, -106.47 |
| ROSE295-08 | D. rosaefolii | MK813666 | Canada | Douglas P.P. | 26-Sep-1999 | 51.04, -106.47 |
| ROSE296-08 | D. rosaefolii | MK813667 | Canada | Douglas P.P. | 26-Sep-1999 | 51.04, -106.47 |
| ROSE303-08 | D. rosaefolii | MK813668 | Canada | Waterton Lakes N.P. | 19-Sep-2007 | 49.1, -113.94 |
| ROSE304-08 | D. rosaefolii | MK813669 | Canada | Waterton Lakes N.P. | 19-Sep-2007 | 49.1, -113.94 |
| ROSE305-08 | D. rosaefolii | MK813670 | Canada | Waterton Lakes N.P. | 19-Sep-2007 | 49.1, -113.94 |
| ROSE306-08 | D. rosaefolii | MK813671 | Canada | Waterton Lakes N.P. | 19-Sep-2007 | 49.1, -113.94 |
| ROSE307-08 | D. rosaefolii | MK813672 | Canada | Waterton Lakes N.P. | 19-Sep-2007 | 49.1, -113.94 |
| ROSE308-08 | D. rosaefolii | MK813673 | Canada | Waterton Lakes N.P. | 19-Sep-2007 | 49.1, -113.94 |
| ROSE309-08 | D. rosaefolii | MK813644 | Canada | Waterton Lakes N.P. | 19-Sep-2007 | 49.1, -113.94 |
| ROSE310-08 | D. rosaefolii | MK813659 | Canada | Waterton Lakes N.P. | 19-Sep-2007 | 49.1, -113.94 |
| ROSE311-08 | D. rosaefolii | MK813657 | Canada | Waterton Lakes N.P. | 19-Sep-2007 | 49.1, -113.94 |
| ROSE312-08 | D. fusiformans | MK813655 | Canada | Waterton Lakes N.P. | 19-Sep-2007 | 49.1, -113.94 |
| ROSE313-08 | D. bassetti | MK813474 | Canada | Coaldale | 24-Oct-2002 | 49.87, -112.43 |
| ROSE314-08 | D. bassetti | MK813473 | Canada | Coaldale | 24-Oct-2002 | 49.87, -112.43 |
| ROSE315-08 | D. bassetti | MK813472 | Canada | Coaldale | 24-Oct-2002 | 49.87, -112.43 |
| ROSE316-08 | D. bassetti | MK813471 | Canada | Coaldale | 24-Oct-2002 | 49.87, -112.43 |
| ROSE317-08 | D. bassetti | MK813470 | Canada | Coaldale | 24-Oct-2002 | 49.87, -112.43 |
| ROSE318-08 | D. bassetti | MK813469 | Canada | Coaldale | 24-Oct-2002 | 49.87, -112.43 |
| ROSE319-08 | D. bassetti | MK813468 | Canada | Coaldale | 24-Oct-2002 | 49.87, -112.43 |
| ROSE320-08 | D. bassetti | MK813467 | Canada | Coaldale | 24-Oct-2002 | 49.87, -112.43 |
| ROSE321-08 | D. bassetti | MK813466 | Canada | Coaldale | 24-Oct-2002 | 49.87, -112.43 |
| ROSE322-08 | D. bassetti | MK813465 | Canada | Osoyoos | 16-Oct-1999 | 49, -119.35 |
| ROSE324-08 | D. bassetti | MK813464 | Canada | Osoyoos | 16-Oct-1999 | 49, -119.35 |
| ROSE325-08 | D. bassetti | MK813463 | Canada | Osoyoos | 16-Oct-1999 | 49, -119.35 |
| ROSE326-08 | D. bassetti | MK813462 | Canada | Osoyoos | 16-Oct-1999 | 49, -119.35 |
| ROSE327-08 | D. bassetti | MK813461 | Canada | Osoyoos | 16-Oct-1999 | 49, -119.35 |
| ROSE328-08 | D. bassetti | MK813460 | Canada | Osoyoos | 16-Oct-1999 | 49, -119.35 |
| ROSE329-08 | D. bassetti | MK813459 | Canada | Osoyoos | 16-Oct-1999 | 49, -119.35 |
| ROSE330-08 | D. bassetti | MK813458 | Canada | Osoyoos | 16-Oct-1999 | 49, -119.35 |
| ROSE332-08 | D. bassetti | MK813457 | Canada | Waterton Lakes N.P. | 19-Sep-2007 | 49.08, -113.9 |
| ROSE333-08 | D. bassetti | MK813456 | Canada | Waterton Lakes N.P. | 19-Sep-2007 | 49.08, -113.9 |
| ROSE334-08 | D. bassetti | MK813479 | Canada | Waterton Lakes N.P. | 19-Sep-2007 | 49.08, -113.9 |
| ROSE335-08 | D. bassetti | MK813478 | Canada | Waterton Lakes N.P. | 19-Sep-2007 | 49.08, -113.9 |
| ROSE336-08 | D. bassetti | MK813477 | Canada | Waterton Lakes N.P. | 19-Sep-2007 | 49.08, -113.9 |
| ROSE337-08 | D. bassetti | MK813476 | Canada | Waterton Lakes N.P. | 19-Sep-2007 | 49.08, -113.9 |
| ROSE338-08 | D. gracilis | MK813554 | Canada | Leader | 08-Sep-1972 | 50.97, -109.53 |
| ROSE341-08 | D. gracilis | MK813555 | Canada | Moose Jaw | 26-Sep-1999 | 51.04, -109.49 |
| ROSE346-08 | D. gracilis | MK813553 | Canada | Saskatoon | 29-Sep-1999 | 51.89, -106.81 |
| ROSE352-08 | D. gracilis | MK813551 | Canada | Waterton Lakes N.P. | 19-Sep-2007 | 49.07, -113.86 |
| ROSE353-08 | D. gracilis | MK813556 | Canada | Waterton Lakes N.P. | 19-Sep-2007 | 49.07, -113.86 |
| ROSE354-08 | D. gracilis | MK813557 | Canada | Waterton Lakes N.P. | 19-Sep-2007 | 49.07, -113.86 |
| ROSE355-08 | D. gracilis | MK813558 | Canada | Waterton Lakes N.P. | 19-Sep-2007 | 49.07, -113.86 |
| ROSE356-08 | D. gracilis | MK813550 | Canada | Waterton Lakes N.P. | 19-Sep-2007 | 49.07, -113.86 |
| ROSE363-08 | D. polita | MK813612 | Canada | Cypress Hills Prov. Park | 20-Oct-1986 | 49.63, -110.23 |
| ROSE365-08 | D. polita | MK813613 | Canada | Cypress Hills Prov. Park | 20-Oct-1986 | 49.63, -110.23 |
| ROSE366-08 | D. polita | MK813614 | Canada | Cypress Hills Prov. Park | 20-Oct-1986 | 49.63, -110.23 |
| ROSE378-08 | D. polita | MK813615 | Canada | Queen Charlotte Islands | 01-Aug-1998 | 52.9, -131.79 |
| ROSE379-08 | D. polita | MK813616 | Canada | Queen Charlotte Islands | 01-Aug-1998 | 52.9, -131.79 |
| ROSE380-08 | D. polita | MK813617 | Canada | Queen Charlotte Islands | 01-Aug-1998 | 52.9, -131.79 |
| ROSE381-08 | D. polita | MK813619 | Canada | Queen Charlotte Islands | 01-Aug-1998 | 52.9, -131.79 |
| ROSE382-08 | D. polita | MK813620 | Canada | Queen Charlotte Islands | 01-Aug-1998 | 52.9, -131.79 |
| ROSE386-08 | D. polita | MK813604 | Canada | Queen Charlotte Islands | 01-Aug-1998 | 52.9, -131.79 |
| ROSE390-08 | D. polita | MK813605 | Canada | Pacific Rim N.P. | 01-Oct-1995 | 49.06, -125.72 |
| ROSE394-08 | D. polita | MK813606 | USA | San Joaquin Co. | 21-Mar-2003 | 37.96, -121.29 |
| ROSE395-08 | D. polita | MK813607 | USA | San Joaquin Co. | 21-Mar-2003 | 37.96, -121.29 |
| ROSE396-08 | D. polita | MK813608 | USA | San Joaquin Co. | 21-Mar-2003 | 37.96, -121.29 |
| ROSE397-08 | D. polita | MK813609 | USA | San Joaquin Co. | 21-Mar-2003 | 37.96, -121.29 |
| ROSE398-08 | D. polita | MK813610 | USA | San Joaquin Co. | 21-Mar-2003 | 37.96, -121.29 |
| ROSE399-08 | D. polita | MK813611 | USA | San Joaquin Co. | 21-Mar-2003 | 37.96, -121.29 |
| ROSE400-08 | D. polita | MK813618 | USA | San Joaquin Co. | 21-Mar-2003 | 37.96, -121.29 |
| ROSE418-08 | D. nodulosa | MK813603 | Canada | Kelowna | 15-Oct-1999 | 49.96, -119.15 |
| ROSE423-08 | D. nebulosa | MK813587 | Canada | Manitoulin Island | 28-Oct-2004 | 45.66, -82.26 |
| ROSE424-08 | D. nebulosa | MK813588 | Canada | Manitoulin Island | 28-Oct-2004 | 45.66, -82.26 |
| ROSE425-08 | D. nebulosa | MK813589 | Canada | Manitoulin Island | 28-Oct-2004 | 45.66, -82.26 |
| ROSE426-08 | D. nebulosa | MK813590 | Canada | Manitoulin Island | 28-Oct-2004 | 45.66, -82.26 |
| ROSE427-08 | D. nebulosa | MK813591 | Canada | Manitoulin Island | 28-Oct-2004 | 45.66, -82.26 |
| ROSE428-08 | D. nebulosa | MK813592 | Canada | Manitoulin Island | 28-Oct-2004 | 45.66, -82.26 |
| ROSE429-08 | D. nebulosa | MK813593 | Canada | Manitoulin Island | 28-Oct-2004 | 45.66, -82.26 |
| ROSE430-08 | D. nebulosa | MK813582 | Canada | Manitoulin Island | 28-Oct-2004 | 45.66, -82.26 |
| ROSE431-08 | D. nebulosa | MK813600 | Canada | Manitoulin Island | 28-Oct-2004 | 45.66, -82.26 |
| ROSE432-08 | D. nebulosa | MK813552 | Canada | Waterton Lakes N.P. | 19-Sep-2007 | 49.1, -113.96 |
| ROSE433-08 | D. nebulosa | MK813599 | Canada | Waterton Lakes N.P. | 19-Sep-2007 | 49.1, -113.96 |
| ROSE434-08 | D. nebulosa | MK813598 | Canada | Waterton Lakes N.P. | 19-Sep-2007 | 49.1, -113.96 |
| ROSE435-08 | D. nebulosa | MK813597 | Canada | Waterton Lakes N.P. | 19-Sep-2007 | 49.1, -113.96 |
| ROSE436-08 | D. nebulosa | MK813596 | Canada | Waterton Lakes N.P. | 19-Sep-2007 | 49.1, -113.96 |
| ROSE437-08 | D. nebulosa | MK813595 | Canada | Waterton Lakes N.P. | 19-Sep-2007 | 49.1, -113.96 |
| ROSE438-08 | D. nebulosa | MK813581 | Canada | Waterton Lakes N.P. | 19-Sep-2007 | 49.1, -113.96 |
| ROSE439-08 | D. nebulosa | MK813602 | Canada | Waterton Lakes N.P. | 19-Sep-2007 | 49.1, -113.96 |
| ROSE440-08 | D. nebulosa | MK813584 | Canada | Waterton Lakes N.P. | 19-Sep-2007 | 49.1, -113.96 |
| ROSE441-08 | D. nebulosa | MK813585 | Canada | Waterton Lakes N.P. | 19-Sep-2007 | 49.1, -113.96 |
| ROSE442-08 | D. nebulosa | MK813546 | Canada | Coaldale | 24-Oct-2002 | 49.87, -112.44 |
| ROSE443-08 | D. nebulosa | MK813547 | Canada | Coaldale | 24-Oct-2002 | 49.87, -112.44 |
| ROSE444-08 | D. nebulosa | MK813583 | Canada | Coaldale | 24-Oct-2002 | 49.87, -112.44 |
| ROSE445-08 | D. nebulosa | MK813580 | Canada | Coaldale | 24-Oct-2002 | 49.87, -112.44 |
| ROSE446-08 | D. nebulosa | MK813579 | Canada | Coaldale | 24-Oct-2002 | 49.87, -112.44 |
| ROSE447-08 | D. nebulosa | MK813578 | Canada | Coaldale | 24-Oct-2002 | 49.87, -112.44 |
| ROSE448-08 | D. nebulosa | MK813577 | Canada | Coaldale | 24-Oct-2002 | 49.87, -112.44 |
| ROSE449-08 | D. nebulosa | MK813548 | Canada | Coaldale | 24-Oct-2002 | 49.87, -112.44 |
| ROSE450-08 | D. nebulosa | MK813601 | Canada | Coaldale | 24-Oct-2002 | 49.87, -112.44 |
| ROSE451-08 | D. nebulosa | MK813594 | Canada | Coaldale | 24-Oct-2002 | 49.87, -112.44 |
| ROSE452-08 | D. fructuum | MK813534 | Turkey | Gümüşhane | 28-Jan-2008 | 40.09, 39.56 |
| ROSE453-08 | D. fructuum | MK813535 | Turkey | Gümüşhane | 28-Jan-2008 | 40.09, 39.56 |
| ROSE454-08 | D. fructuum | MK813516 | Turkey | Gümüşhane | 28-Jan-2008 | 40.09, 39.56 |
| ROSE455-08 | D. fructuum | MK813521 | Turkey | Gümüşhane | 28-Jan-2008 | 40.09, 39.56 |
| ROSE456-08 | D. fructuum | MK813532 | Turkey | Gümüşhane | 28-Jan-2008 | 40.09, 39.56 |
| ROSE457-08 | D. fructuum | MK813517 | Turkey | Gümüşhane | 28-Jan-2008 | 40.09, 39.56 |
| ROSE458-08 | D. fructuum | MK813518 | Turkey | Gümüşhane | 28-Jan-2008 | 40.09, 39.56 |
| ROSE459-08 | D. fructuum | MK813519 | Turkey | Gümüşhane | 28-Jan-2008 | 40.09, 39.56 |
| ROSE460-08 | D. fructuum | MK813520 | Turkey | Gümüşhane | 28-Jan-2008 | 40.09, 39.56 |
| ROSE461-08 | D. fructuum | MK813522 | Turkey | Erzurum | 12-Jan-2008 | 39.9, 41.24 |
| ROSE462-08 | D. fructuum | MK813523 | Turkey | Erzurum | 12-Jan-2008 | 39.9, 41.24 |
| ROSE463-08 | D. fructuum | MK813524 | Turkey | Erzurum | 12-Jan-2008 | 39.9, 41.24 |
| ROSE464-08 | D. fructuum | MK813525 | Turkey | Erzurum | 12-Jan-2008 | 39.9, 41.24 |
| ROSE465-08 | D. fructuum | MK813526 | Turkey | Erzurum | 12-Jan-2008 | 39.9, 41.24 |
| ROSE466-08 | D. fructuum | MK813527 | Turkey | Erzurum | 12-Jan-2008 | 39.9, 41.24 |
| ROSE467-08 | D. fructuum | MK813528 | Turkey | Erzurum | 12-Jan-2008 | 39.9, 41.24 |
| ROSE468-08 | D. fructuum | MK813529 | Turkey | Erzurum | 12-Jan-2008 | 39.9, 41.24 |
| ROSE469-08 | D. fructuum | MK813530 | Turkey | Erzurum | 12-Jan-2008 | 39.9, 41.24 |
| ROSE470-08 | D. fructuum | MK813531 | Turkey | Erzurum | 12-Jan-2008 | 39.9, 41.24 |
| ROSE488-08 | D. rosae | MK813631 | Canada | St. John's | 10-Oct-1987 | 47.58, -52.69 |
| ROSE489-08 | D. rosae | MK813630 | Canada | St. John's | 10-Oct-1987 | 47.58, -52.69 |
| ROSE490-08 | D. rosae | MK813641 | Canada | St. John's | 10-Oct-1987 | 47.58, -52.69 |
| ROSE491-08 | D. rosae | MK813640 | Canada | St. John's | 10-Oct-1987 | 47.58, -52.69 |
| ROSE492-08 | D. rosae | MK813639 | Canada | St. John's | 10-Oct-1987 | 47.58, -52.69 |
| ROSE493-08 | D. rosae | MK813638 | Canada | Prince Edward Co. | 28-Apr-2003 | 43.95, -76.87 |
| ROSE494-08 | D. rosae | MK813637 | Canada | Prince Edward Co. | 28-Apr-2003 | 43.95, -76.87 |
| ROSE495-08 | D. rosae | MK813636 | Canada | Prince Edward Co. | 28-Apr-2003 | 43.95, -76.87 |
| ROSE496-08 | D. rosae | MK813635 | Canada | Prince Edward Co. | 28-Apr-2003 | 43.95, -76.87 |
| ROSE497-08 | D. rosae | MK813633 | Canada | Malden Center | 01-May-1994 | 42.06, -83.06 |
| ROSE498-08 | D. rosae | MK813643 | Canada | Malden Center | 01-May-1994 | 42.06, -83.06 |
| ROSE499-08 | D. rosae | MK813629 | Canada | Malden Center | 01-May-1994 | 42.06, -83.06 |
| ROSE500-08 | D. rosae | MK813634 | Canada | Malden Center | 01-May-1994 | 42.06, -83.06 |
| ROSE501-08 | D. rosae | MK813642 | Canada | Malden Center | 01-May-1994 | 42.06, -83.06 |
| ROSE502-08 | D. eglanteriae | MK813514 | Canada | Niagara Falls | 06-May-1996 | 43.09, -79.09 |
| ROSE504-08 | D. eglanteriae | MK813513 | Canada | Niagara Falls | 06-May-1996 | 43.09, -79.09 |
| ROSE505-08 | D. eglanteriae | MK813512 | Canada | Niagara Falls | 06-May-1996 | 43.09, -79.09 |
| ROSE506-08 | D. eglanteriae | MK813507 | Canada | Niagara Falls | 06-May-1996 | 43.09, -79.09 |
| ROSE515-08 | D. californica | MK813500 | USA | San Diego Co. | 17-Aug-1969 | 33.34, -116.91 |
| ROSE516-08 | D. californica | MK813499 | USA | San Diego Co. | 17-Aug-1969 | 33.34, -116.91 |
| ROSE518-08 | D. californica | MK813501 | USA | San Diego Co. | 17-Aug-1969 | 33.34, -116.91 |
| ROSE519-08 | D. californica | MK813502 | USA | Davis | 24-Feb-1993 | 38.35, -122.27 |
| ROSE520-08 | D. californica | MK813503 | USA | Davis | 24-Feb-1993 | 38.35, -122.27 |
| ROSE521-08 | D. californica | MK813504 | USA | Davis | 24-Feb-1993 | 38.35, -122.27 |
| ROSE523-08 | D. californica | MK813505 | USA | Davis | 24-Feb-1993 | 38.35, -122.27 |
| ROSE524-08 | D. californica | MK813506 | USA | Davis | 24-Feb-1993 | 38.35, -122.27 |
| ROSE553-08 | D. japonica | MK813441 | Japan | Kuroishi | 01-Nov-1996 | 40.64, 140.6 |
| ROSE554-08 | D. japonica | MK813440 | Japan | Kuroishi | 01-Nov-1996 | 40.81, 140.76 |
| ROSE565-08 | D. spinosa | MK813688 | Canada | Waterton Lakes N.P. | 10-May-2007 | 49.08, -113.88 |
| ROSE566-08 | D. spinosa | MK813687 | Canada | Waterton Lakes N.P. | 10-May-2007 | 49.08, -113.88 |
| ROSE567-08 | D. spinosa | MK813686 | Canada | Waterton Lakes N.P. | 10-May-2007 | 49.08, -113.88 |
| ROSE568-08 | P. pirata | MK813827 | Canada | Waterton Lakes N.P. | 10-May-2007 | 49.08, -113.88 |
| ROSE569-08 | P. pirata | MK813826 | Canada | Waterton Lakes N.P. | 10-May-2007 | 49.08, -113.88 |
| ROSE570-08 | P. pirata | MK813829 | Canada | Waterton Lakes N.P. | 10-May-2007 | 49.08, -113.88 |
| ROSE571-08 | P. pirata | MK813828 | Canada | Waterton Lakes N.P. | 10-May-2007 | 49.08, -113.88 |
| ROSE572-08 | P. pirata | MK813830 | Canada | Waterton Lakes N.P. | 10-May-2007 | 49.08, -113.88 |
| ROSE609-08 | D. spinosa | MK813685 | Canada | Waterton Lakes N.P. | 10-May-2007 | 49.1, -113.96 |
| ROSE610-08 | D. spinosa | MK813684 | Canada | Waterton Lakes N.P. | 10-May-2007 | 49.1, -113.96 |
| ROSE611-08 | D. spinosa | MK813683 | Canada | Waterton Lakes N.P. | 10-May-2007 | 49.1, -113.96 |
| ROSE612-08 | P. pirata | MK813831 | Canada | Waterton Lakes N.P. | 10-May-2007 | 49.08, -113.88 |
| ROSE659-09 | D. nodulosa | GU695239 | Canada | Manitoulin Island | 30-Apr-2002 | 45.83, -82.33 |
| ROSE660-09 | D. nodulosa | GU695240 | Canada | Manitoulin Island | 30-Apr-2002 | 45.83, -82.33 |
| ROSE661-09 | D. nodulosa | GU695241 | Canada | Water Lakes N.P. | 24-Sep-2007 | 49.08, -113.87 |
| ROSE662-09 | D. nodulosa | GU695235 | Canada | Manitoulin Island | 28-Apr-2000 | 45.66, -82.26 |
| ROSE663-09 | D. nodulosa | GU695236 | Canada | Manitoulin Island | 28-Apr-2000 | 45.66, -82.26 |
| ROSE664-09 | D. nodulosa | GU695237 | Canada | Manitoulin Island | 28-Apr-2000 | 45.66, -82.26 |
| ROSE665-09 | D. nodulosa | GU695238 | Canada | Manitoulin Island | 28-Apr-2000 | 45.66, -82.26 |
| ROSE666-09 | D. nodulosa | GU695231 | Canada | Manitoulin Island | 28-Apr-2000 | 45.66, -82.26 |
| ROSE667-09 | D. nodulosa | GU695232 | Canada | Manitoulin Island | 26-Aug-1999 | 45.66, -82.26 |
| ROSE668-09 | D. nodulosa | GU695233 | Canada | Manitoulin Island | 26-Aug-1999 | 45.66, -82.26 |
| ROSE691-09 | D. radicum | GU695213 | Canada | Fort Albany | 25-May-2005 | 52.25, -81.5 |
| ROSE693-09 | D. radicum | GU695214 | Canada | Fort Albany | 25-May-2005 | 52.25, -81.5 |
| ROSE694-09 | D. radicum | GU695215 | Canada | Fort Albany | 25-May-2005 | 52.25, -81.5 |
| ROSE696-09 | D. radicum | GU695209 | Canada | Fort Albany | 25-May-2005 | 52.25, -81.5 |
| ROSE698-09 | D. radicum | GU695210 | Canada | Fort Albany | 25-May-2005 | 52.25, -81.5 |
| ROSE701-09 | D. oregonensis | GU695211 | Canada | Sceptre | 09-May-2003 | 50.44, -109.22 |
| ROSE702-09 | D. oregonensis | GU695212 | Canada | Sceptre | 09-May-2003 | 50.44, -109.22 |
| ROSE704-09 | D. oregonensis | GU695205 | Canada | Sceptre | 09-May-2003 | 50.44, -109.22 |
| ROSE705-09 | D. oregonensis | GU695206 | Canada | Sceptre | 09-May-2003 | 50.44, -109.22 |
| ROSE706-09 | D. oregonensis | GU695207 | Canada | Sceptre | 09-May-2003 | 50.44, -109.22 |
| WSU14 | D. oregonensis | MK977681 | USA | Pullman | 01-May-2007 | 46.74, -117.14 |
| WSUHip17 | D. oregonensis | MK977682 | USA | Pullman | 01-May-2007 | 46.74, -117.14 |
| WSUPP03 | D. oregonensis | MK977680 | USA | Pullman | 01-May-2007 | 46.74, -117.14 |

**Supplementary Table 2.** Intraspecific genetic divergence of A) *Diplolepis* and B) *Periclistus*.

| **A** |  |  |
| --- | --- | --- |
| **Species** | **Intraspecific Divergence (%)** | **Standard Deviation** |
| *D. eglanteriae* Palearctic | 2.44 | 0.58 |
| *D. rosae* Group | 1.86 | 0.37 |
| *Diplolepis* sp.2 | 0.12 | 0.11 |
| *D. spinosissimae* | 1.65 | 0.40 |
| *D. nervosa* | 0.35 | 0.20 |
| *D. oregonensis* | 1.66 | 0.35 |
| *D. bicolor* | 1.46 | 0.32 |
| *D. californica* | 1.10 | 0.26 |
| *D. rosaefolii* Group | 2.48 | 0.43 |
| *D. ignota* Group | 0.45 | 0.17 |
| *D. triforma* | 0.00 | 0.00 |
| *D. spinosa* | 1.76 | 0.36 |
| *D. polita* Group | 2.41 | 0.40 |
| *D. radicum* | 1.44 | 0.22 |
| *D. eglanteriae* Nearctic | 0.00 | 0.00 |
| *D. gracilis* | 0.04 | 0.04 |
| *D. nodulosa* | 0.23 | 0.10 |
| **B** |  |  |
| **Species** | **Intraspecific Divergence (%)** | **Standard Deviation** |
| *P. brandtii* | 0.28 | 0.22 |
| *P. caninae* | 1.46 | 0.60 |
| *P. arefactus* | 0.13 | 0.14 |
| *Periclistus* sp.1 | 2.16 | 0.66 |
| *P. piceus* | 0.64 | 0.22 |
| *P. pirata* | 1.47 | 0.49 |
| *Periclistus* sp.2 | 0.30 | 0.17 |

**Supplementary Table 3.** Interspecific genetic divergence of A) *Diplolepis* and B) *Periclistus*.

| **A** |  |  |  |  |  |  |  |  |  |  |  |  |  |  |  |  |  |
| --- | --- | --- | --- | --- | --- | --- | --- | --- | --- | --- | --- | --- | --- | --- | --- | --- | --- |
| **Species** | *D.eglanteriae* Palearctic | *D.rosae* Group | *Diplolepis* sp.2 | *D. spinosissimae* | *D. nervosa* | *D. oregonensis* | *D. bicolor* | *D. californica* | *D. rosaefolii* Group | *D. ignota* Group | *D. triforma* | *D. spinosa* | *D. polita* Group | *D. radicum* | *D. eglanteriae* Nearctic | *D. gracilis* | *D. nodulosa* |
| *D.eglanteriae* Palearctic |  | 1.6 | 1.68 | 1.65 | 1.36 | 1.64 | 1.46 | 1.81 | 1.55 | 1.60 | 1.78 | 1.77 | 1.41 | 1.85 | 1.08 | 1.60 | 1.81 |
| *D.rosae* Group | 13.68 |  | 0.97 | 0.99 | 1.43 | 1.23 | 1.27 | 1.45 | 1.11 | 1.18 | 1.43 | 1.52 | 1.21 | 1.41 | 1.63 | 1.25 | 1.41 |
| *Diplolepis* sp.2 | 14.20 | 5.93 |  | 0.89 | 1.52 | 1.40 | 1.30 | 1.60 | 1.34 | 1.21 | 1.46 | 1.58 | 1.26 | 1.53 | 1.77 | 1.21 | 1.38 |
| *D. spinosissimae* | 14.17 | 6.28 | 4.60 |  | 1.32 | 1.52 | 1.23 | 1.61 | 1.22 | 1.19 | 1.51 | 1.62 | 1.13 | 1.57 | 1.64 | 1.34 | 1.47 |
| *D. nervosa* | 11.31 | 10.94 | 11.83 | 9.94 |  | 1.67 | 1.48 | 1.64 | 1.38 | 1.48 | 1.62 | 1.72 | 1.44 | 1.71 | 1.45 | 1.56 | 1.67 |
| *D. oregonensis* | 14.33 | 10.13 | 11.36 | 12.64 | 14.28 |  | 1.46 | 1.10 | 1.34 | 1.33 | 1.13 | 1.08 | 1.46 | 1.02 | 1.66 | 1.36 | 1.15 |
| *D. bicolor* | 12.87 | 10.24 | 10.29 | 9.33 | 11.90 | 12.67 |  | 1.75 | 1.39 | 1.39 | 1.63 | 1.72 | 0.84 | 1.67 | 1.55 | 1.44 | 1.67 |
| *D. californica* | 17.20 | 11.89 | 13.05 | 13.23 | 13.79 | 8.05 | 16.43 |  | 1.50 | 1.54 | 1.06 | 0.99 | 1.67 | 0.90 | 1.73 | 1.48 | 1.17 |
| *D. rosaefolii* Group | 13.74 | 8.92 | 10.43 | 9.11 | 10.51 | 12.00 | 11.22 | 12.85 |  | 1.27 | 1.38 | 1.43 | 1.30 | 1.47 | 1.65 | 1.30 | 1.43 |
| *D. ignota* Group | 13.97 | 8.98 | 8.51 | 8.20 | 11.87 | 10.45 | 11.49 | 13.09 | 10.39 |  | 1.42 | 1.50 | 1.43 | 1.51 | 1.56 | 0.94 | 1.45 |
| *D. triforma* | 17.01 | 11.52 | 11.75 | 12.25 | 13.54 | 8.37 | 15.14 | 6.95 | 12.31 | 11.29 |  | 1.00 | 1.56 | 1.05 | 1.68 | 1.50 | 0.97 |
| *D. spinosa* | 17.03 | 13.73 | 13.12 | 14.14 | 14.69 | 8.97 | 16.34 | 7.35 | 12.85 | 12.82 | 7.10 |  | 1.69 | 0.94 | 1.71 | 1.51 | 1.11 |
| *D. polita* Group | 12.99 | 9.59 | 9.89 | 8.73 | 11.88 | 12.83 | 6.08 | 15.61 | 10.40 | 12.16 | 14.06 | 15.97 |  | 1.65 | 1.54 | 1.36 | 1.61 |
| *D. radicum* | 17.76 | 12.16 | 12.76 | 13.36 | 14.70 | 6.91 | 15.89 | 6.09 | 13.44 | 12.77 | 7.03 | 6.92 | 15.56 |  | 1.81 | 1.57 | 1.15 |
| *D. eglanteriae* Nearctic | 7.16 | 13.73 | 14.61 | 13.75 | 10.90 | 14.18 | 13.58 | 15.78 | 14.05 | 12.80 | 15.21 | 15.73 | 13.97 | 16.79 |  | 1.54 | 1.77 |
| *D. gracilis* | 13.30 | 10.09 | 9.41 | 10.17 | 13.56 | 11.60 | 11.66 | 12.57 | 10.00 | 5.76 | 12.60 | 13.51 | 11.34 | 14.31 | 13.80 |  | 1.49 |
| *D. nodulosa* | 17.55 | 11.02 | 10.44 | 11.60 | 14.15 | 8.43 | 15.23 | 7.64 | 12.07 | 11.86 | 5.99 | 7.98 | 14.51 | 7.66 | 16.71 | 12.36 |  |
| **B** |  |  |  |  |  |  |  |  |  |  |  |  |  |  |  |  |  |
| **Species** | *P. brandtii* | *P. caninae* | *P. arefactus* | *Periclistus* sp.1 | *P. piceus* | *P. pirata* | *Periclistus* sp.2 |  |  |  |  |  |  |  |  |  |  |
| *P. brandtii* |  | 1.33 | 2.40 | 2.86 | 2.41 | 2.92 | 2.92 |  |  |  |  |  |  |  |  |  |  |
| *P. caninae* | 4.28 |  | 2.26 | 2.44 | 2.00 | 2.53 | 2.42 |  |  |  |  |  |  |  |  |  |  |
| *P. arefactus* | 7.68 | 7.40 |  | 2.44 | 2.20 | 1.10 | 2.40 |  |  |  |  |  |  |  |  |  |  |
| *Periclistus* sp.1 | 9.05 | 7.92 | 7.96 |  | 2.53 | 2.74 | 1.14 |  |  |  |  |  |  |  |  |  |  |
| *P. piceus* | 7.87 | 6.57 | 7.09 | 7.96 |  | 2.33 | 2.69 |  |  |  |  |  |  |  |  |  |  |
| *P. pirata* | 9.30 | 8.23 | 3.55 | 8.92 | 7.64 |  | 2.74 |  |  |  |  |  |  |  |  |  |  |
| *Periclistus* sp.2 | 9.16 | 7.84 | 7.79 | 3.67 | 8.38 | 8.86 |  |  |  |  |  |  |  |  |  |  |  |
